# The trypanosomatid dynamin-like protein associates with glycosomes

**DOI:** 10.64898/2026.04.27.721030

**Authors:** Madeline F. Malfara, Blyssalyn V. Bieber, Rodolpho O. O. Souza, Thomas Beer, Hsin-Yao Tang, Megan L. Povelones

**Affiliations:** Department of Biology, Villanova University, Villanova, PA USA; Department of Microbiology and Immunology, University at Buffalo, Buffalo, NY USA; Wistar Institute, Philadelphia, PA USA

## Abstract

Subcellular organelles must undergo periodic fission to be evenly distributed during cell division. These division events are mediated by protein members of the dynamin family, including dynamin-related proteins. Protozoan parasites, including trypanosomatids such as *Trypanosoma brucei*, have several single-copy organelles, suggesting tightly regulated systems for organelle fission and segregation. However, trypanosomatid genomes typically encode only one dynamin-like protein (DLP), which in *T. brucei* has multiple roles including endocytosis and mitochondrial fission. How DLPs are recruited to different membranes, and how their fission activity is regulated, are unknown. We used tandem-affinity purification in the related trypanosomatid *Crithidia fasciculata* to identify interacting partners of DLP. Surprisingly, we found that *Cf*DLP co-purified with multiple proteins predicted to localize to glycosomes, peroxisome-related glycolytic organelles. Using expansion microscopy, we confirmed the localization of *Cf*DLP to glycosomes, specifically those that appear to be undergoing division. To see if changes in the levels of DLP could alter glycosome morphology, we conducted RNAi-mediated knockdown and inducible overexpression experiments in *T. brucei*. *Tb*DLP knockdown causes subtle changes in glycosome size, while overexpression of *Tb*DLP1 causes an increase cytoplasmic vesicles and altered permeability of glycosomal membranes. These results suggest that the multifunctional DLP of trypanosomatids plays a role in glycosome maintenance.

**Author Summary:** Trypanosomatids are eukaryotic parasites that cause devastating diseases in humans and animals. Like all eukaryotic cells, they must maintain their subcellular compartments through organelle division and other membrane remodeling events. Dynamin-like proteins are enzymes that work with other proteins to apply mechanical force to membranes. The dynamin-like proteins of *Trypanosoma brucei*, the causative agent of human African trypanosomiasis, have been implicated in endocytosis and mitochondrial division, although how these activities are regulated is not known. We have used a model trypanosomatid, the mosquito parasite *Crithidia fasciculata*, to look for dynamin-interacting proteins. In addition to proteins of unknown function, we show that dynamin-like protein associates with proteins found on glycosomes, trypanosomatid-specific organelles that contain enzymes required for breakdown of sugars. Knockdown and overexpression of dynamin-like proteins in *T. brucei* causes changes in glycosomes, supporting a role in organelle maintenance. Dynamin-like proteins likely regulate organelle structure and function, allowing parasites to adapt to different energetic requirements during their life cycle.

## Introduction

Trypanosomatids are single-celled eukaryotic parasites, several of which cause important diseases in humans and livestock (World Health Organization, 2025; World Organization for Animal Health (2025), 2025). They have a distinctly polarized cellular organization required for survival in both their mammalian hosts and the insect vectors that transmit them (3). In this regard, the parasites face two main challenges. First, they possess several single-copy organelles, such as the flagellum, mitochondrion, mitochondrial nucleoid (kinetoplast DNA or kDNA), and Golgi body (4,5), which must be faithfully replicated and segregated during each cell cycle. Second, the structure of their energy-producing organelles changes during the developmental cycle of the parasite to adapt to different nutrient environments (6–8). This includes remodeling of the mitochondrion and glycosomes, peroxisome-like organelles that sequester the first seven steps of glycolysis as well as other metabolic pathways (9). Glycosomes also play a role in signal transduction during differentiation (10,11). Unlike the single mitochondrion, trypanosomatids contain many glycosomes per cell which are heterogenous in composition (12,13).

The single flagellum emerges from an invagination called the flagellar pocket (14). In *Trypanosoma brucei* and *Leishmania*, all cellular endo- and exocytosis takes place at this specialized membrane (15). Long slender bloodstream form (BSF) *T. brucei* has a rapid endocytic rate, allowing these obligate extracellular parasites to shed immune effectors from its surface (16). Endocytosis in procyclic form (PCF) *T. brucei*, found in the midgut of the tsetse fly vector, still occurs at the flagellar pocket, but at a 10-fold lower rate (17).

In traditional model eukaryotes, scission of endocytic vesicles is mediated by dynamins, a conserved family of GTPases that perform membrane remodeling, usually through formation of ring-like oligomers (18). Classical dynamins interact with the membrane via pleckstrin-homology (PH) domains. Dynamin-related or dynamin-like proteins (DLPs) lack PH domains and interact with adaptor proteins in organelle membranes to mediate organelle fission (19).

While yeast and metazoa have both classical dynamins and DLPs, early-diverging eukaryotes such as trypanosomatids and apicomplexan parasites have only DLPs. In *T. gondii*, *Tg*DrpB is involved in biogenesis of secretary organelles (20) while *Tg*DrpC has been linked to mitochondrial fission (21) as well as vesicle transport and cell division (22,23). *Tg*DrpA plays a role in division of the apicoplast, a non-photosynthetic plastid, localizing to the point of apicoplast fission during endodyogeny (24) and suggesting formation of a dynamin ring similar to mitochondrial fission machinery (25). In *P. falciparum*, *Pf*Dyn1 may be important for vesicle budding during uptake of hemoglobin (26) and *Pf*Dyn2 has been shown to be required for fission of both the mitochondrion and the apicoplast (27–30).

*T. brucei* has two, nearly identical dynamin-like proteins called *Tb*DLP1 and *Tb*DLP2 (31–33), as well as a more distantly related dynamin family member called *Tb*MFNL. Overexpression of *Tb*MFNL changes mitochondrial structure in a GTPase-dependent manner, but is from a lineage distinct from DLP and causes no apparent phenotype when knocked out (34,35). Although they differ by only 19 amino acids, *Tb*DLP1 and *Tb*DLP2 are differentially regulated during the cell and life cycle of the parasite and play overlapping yet distinct roles in endocytosis and mitochondrial fission (31). Interestingly, related parasites in the *Leishmania* lineage, including monoxenous parasites of insects such as *Crithidia fasciculata*, have only a single DLP. In *L. donovani*, this protein has been shown to self-oligomerize (36) and mutations have been linked to drug resistance (37). How trypanosomatid DLPs are recruited to different internal membranes and why *T. brucei* requires two DLPs are not understood. In addition, since each cell has a single mitochondrion, fission of this organelle must be timed with the cell cycle, and may be mechanistically linked to cytokinesis (32,38). *T. brucei* and *C. fasciculata* lack orthologs to most adaptor proteins present in yeast and mammalian cells, and to date no adaptors have been functionally characterized in these organisms. To better understand the role of trypanosomatid DLP in organelle division and endocytosis, we sought to identify putative DLP interactors through tandem-affinity purification and mass spectrometry in *C. fasciculata*. We present evidence that DLP in trypanosomatids associates with glycosomes, identifying this protein as a master regulator of cellular organization and function in trypanosomatids.

## Methods

### Cell culture

*Crithidia fasciculata* strain CfC1 (39) was cultured in brain heart infusion (BHI) medium supplemented with hemin at 27 °C as described (40). Ectopic plasmids were maintained using 200 µg/mL hygromycin. Procyclic form *Trypanosoma brucei brucei* strain 29-13 (41,42) were maintained at 27 °C in SDM-79 medium supplemented with 15% fetal calf serum (Atlanta Biologicals) and 15 µg/mL neomycin (G418, Millipore Sigma), 50 µg/mL hygromycin (ThermoFisher), 5 µg/mL blasticidin (ThermoFisher), 0.1 µg/mL puromycin (ThermoFisher), and 2.5µg/mL phleomycin (Millipore Sigma) as appropriate.

### Plasmids

*Cf*DLP (CFAC1_200029900) was amplified by PCR (see Table S1) and cloned into pNUS-PTPcH (43) using NdeI and NotI (an additional nucleotide was added to the reverse primer to restore the reading frame between *Cf*DLP and the PTP tag) to create pNUS-PTPcH-CfDLP. pNUS-PTPcH-CfRfc3 and pNUS-PTPcH-CfRrp4 were a gift from Stuart MacNeill (43). The full open reading frame of the putative *C. fasciculata* Gim5A (CFAC1_300087400) was amplified by PCR and cloned using NdeI and SphI into the pNUS-HcH vector that had been previously modified to include a single myc tag before the 6X His tag (44). Constructs were introduced as episomes into *C. fasciculata* strain CfC1 using a Lonza Nucleofector IIb as described (38). After seven days of selection with 80 µg/mL hygromycin, the drug concentration was increased to 200 µg/mL. Experiments with PTP cell lines were performed on populations. For pNUS-MYCcH-*Cf*Gim5A, selected cells were cloned by limiting dilution in a 96-well plate and screened by immunofluorescence. The size of the tagged protein was confirmed by western blot with anti-c-myc monoclonal antibody (9E10.3) and the localization of the tagged protein was compared to that of the polyclonal aldolase antibody (a gift of Meredith Morris). The pXS2aldoPTS2eYFP construct was also provided by Meredith Morris, and contains the PTS2 signal from *T. brucei* aldolase cloned into pXS2 and fused by its N-terminus to eYFP as described (45). A stem-loop RNAi construct targeting *Tb*DLP1/2 was provided by Andre Schneider (32). *Tb*DLP1 was PCR amplified with a C-terminal Ty1 tag included on the reverse primer and cloned into pLEW100v5 (a gift from George Cross, Addgene plasmid #24011) using Gibson Assembly (NEB). For each of these constructs, 5-10 µg of linearized plasmid was transfected into *T. brucei* PCF cells using Tb-BSF buffer (90 mM sodium phosphate, 5 mM potassium chloride, 0.15 mM calcium chloride, 50 mM HEPES, pH 7.3) (46) and a Lonza IIb Nucelofector set to program X-014. The pXSaldoPTS2eYFP construct was introduced into strain 29-13, the *Tb*DLP1/2 stem-loop RNAi construct was introduced into the TbPCF-mitoGFP background (47), and the TbDLP1::Ty overexpression construct was introduced into a cell line containing an mNeonGreen-tagged *Tb*VAP protein created as described (48). Cells were selected with either blasticidin, puromycin, or phleomycin and clonal lines were obtained through limiting dilution.

### Structural predictions

Structure predictions were made using a modified version of Colabfold Alphafold2 iPython notebook optimized to improve protein predictions for species in the Discoba lineage (**49–51**). Protein sequences were obtained from TriTrypDB (52). Molecular graphics highlighting particular residues were created in UCSF ChimeraX, developed by the Resource for Biocomputing, Visualization, and Informatics at the University of California, San Francisco, with support from National Institutes of Health R01-GM129325 and the Office of Cyber Infrastructure and Computational Biology, National Institute of Allergy and Infectious Diseases (**53–55**). *C. fasciculata* DLP putative phosphosites S260, S503, S592 and T594 were identified in mass spectrometry experiments. *T. brucei* DLP1 phosphosites S537 and S540 were previously identified (56,57).

### Tandem affinity purification

*Cf*DLP-PTP and *Cf*Rrp4-PTP were purified as in (43,58). 100 mL of cells were grown to mid-log (∼5E7 cells/mL) density in BHI and harvested by centrifugation for 10 minutes. Cells were washed 3 times in 5 mL ice-cold 1x PBS and resuspended in 1.5 volume ice-cold PA-300 (150 mM sucrose, 300 mM potassium chloride, 40 mM potassium L-glutamate, 3 mM MgCl_2_. 20 mM HEPES KOH (pH 7.7), 2 mM DTT, 0.1% Tween-20, 1x complete mini EDTA-free protease inhibitor cocktail from Roche). Cells were lysed in a 7 mL Dounce homogenizer (Millipore Sigma) using continuous strokes for 5 minutes on ice in a cold room. Following lysis, cells were centrifuged at 20,500 xg for 10 minutes at 4 °C. Cell lysate was filtered into a 10 mL poly-prep chromatography column (Bio-Rad), that was pre-equilibrated with 200 µL of IgG Sepharsoe 6 Fastflow beads (GE Healthcare) and PA-300 buffer, and rotated for 2 hours at 4°C. Beads were washed twice with PA-300 buffer and equilibrated with TEV buffer (150 mM potassium chloride, 20 mM Tris-HCl (pH 7.7), 3 mM MgCl_2_, 0.5 mM DTT, 0,1% Tween-20, 0.5 mM EDTA) before addition of 200 U of AcTEV protease (ThermoFisher) followed by an overnight incubation at 4 °C. The next day the eluate was added to a second poly-Prep column pre-equilibrated with 200 µL of anti-ProtC Affinity Matrix beads in PC-150 buffer (150 mM potassium chloride, 20 mM Tris-HCl pH 7.7, 3 mM MgCl_2_, 0.5 mM DTT, 0,1% Tween-20, 1mM CaCl_2_, 1x complete mini EDTA-free protease inhibitor cocktail) and rotated for 2 hours at 4 °C. Columns were washed 6 times with PC-150 buffer and samples were eluted in 1.8 mL of room temperature EGTA/EDTA buffer (5 mM Tris-HCl pH 7.7, 10 mM EGTA, 5 mM EDTTA, 10 µg/mL leupeptin) and concentrated using 30 µL of StrataClean Resin (Agilent) before boiling in 20 µL of 4x NuPAGE LDS sample buffer (Thermo Fisher Scientific). 20 µL of purified protein sample was run on a pre-cast SDS-PAGE gel and stained with SYPRO-Ruby prior to excision, trypsinization and analysis by mass spectrometry.

### Mass Spectrometry and Proteomic Analysis

To identify proteins interacting with *Cf*DLP, we performed liquid chromatography-tandem mass spectrometry (LC-MS/MS) on triplicate samples of *Cf*DLP::PTP and duplicate samples of *Cf*RRP4::PTP (control). Following purification, samples were resolved 1 cm into a 10% non-reducing NuPage SDS-PAGE gel (Thermofisher) and visualized with Colloidal Blue staining. The entire stained gel regions were excised and subjected to in-gel trypsin digestion after reduction with TCEP and alkylation with iodoacetamide. Tryptic peptides were analyzed using a 95-min LC method on a Thermo Q Exactive Plus mass spectrometer interfaced with a NanoAcquity UPLC system (Waters) as described previously (59). MS data were searched against the *C. fasciculata* proteome database (TriTrypDB-9.0_CfasciculataCfCl_AnnotatedProteins) using MaxQuant v1.6.17.0 (60) with full tryptic specificity. Carbamidomethylation of Cys was set as a fixed modification. Variable modifications considered in the search were protein N-terminal acetylation, Asn deamidation, and Met oxidation. A separate search incorporating Ser, Thr and Tyr phosphorylation as variable modifications was also performed to identify phosphorylation events. Peptide, protein, and site false discovery rates (FDRs) were set to 1% using a target-decoy reverse database. For downstream analysis, protein intensities were Log_2_-transformed, and missing values were imputed with a minimum value to facilitate statistical comparisons. Differentially abundant proteins were defined by a Log_2_ fold-change (DLP/RRP4) >2 and a p-value <0.05. The mass spectrometry proteomics data have been deposited in the ProteomeXchange Consortium through the MassIVE repository under the dataset identifier PXD074777 and MSV000100956.

### Ultrastructure Expansion Microscopy (U-ExM)

Ultrastructure Expansion Microscopy (U-ExM) was performed as described previously (61,62) with the following modifications: parasites were fixed in 2% paraformaldehyde and placed on poly-D-lysine-coated coverslips. Following fixation, cells were washed twice in 1X PBS+0.1M glycine and stored in 1X PBS until further processing for expansion microscopy. Primary antibodies were mouse anti-GFP (168AT1211 cat. no. AM1009A, ABGENT, 1:250 dilution), mouse anti-cMyc (clone 9E10, Cell Signaling, 1:500 dilution), and rabbit anti-DLP [Boster Bio, affinity-purified antibody raised against AA 1-227 of Tb927.3.4760 (*Tb*DLP2); 1:500 dilution]. Secondary antibodies were Alexa Fluor NHS 405 NHS-ester, Alexa Fluor 647-, Alexa Fluor 594-, or Alexa Fluor 488-conjugated goat anti-rabbit-IgG and goat anti-mouse-IgG (Invitrogen). Secondary antibodies were used at 1:1000 except for the Alexa Fluor NHS-ester 405, which was used at 1:250. Images were acquired on an LSM 900 microscope using a 63× Plan-Apochromat (NA 1.4) objective lens as Z-stacks with an XY pixel size of 0.035 µm and a Z-step size of 0.15 µm. Images were processed with Airyscan using ZEN Blue (Version 3.1, Zeiss, Oberkochen, Germany). Images were later processed and analyzed using FIJI ImageJ 64 Software (63). After the second round of expansion, the gels were measured with a ruler, and the expansion factor (the gel size relative to the coverslip) was calculated. The expansion factor for this experiment was 4.5x. The TurboReg FIJI plugin was used to adjust registration of images shifted during z-stack (settings: translation, manual, accurate) (64,65).

### Fluorescence microscopy

Either 1E7 *T. brucei* or 5E6 *C. fasciculata* cells from mid-log cultures were centrifuged at 800 xg for 5 minutes, washed once in 1X PBS and added to glass coverslips pretreated with 0.1% poly-L-lysine. Cells were allowed to adhere to coverslips for 10 minutes (*T. brucei)* or 20 minutes (*C. fasciculata*) at room temperature in a humid chamber. Cells were washed with PBS and fixed in cold 4% PFA for 15 minutes at room temperature. Following fixation, coverslips were washed in PBS containing 0.1M glycine and permeabilized for 5 minutes in 1% Triton X-100. Cells were stained with primary antibody (Rabbit anti-DLP, Boster Bio 1:500; Rabbit anti-aldolase, 1:500; Mouse anti-c-myc 9E10, ThermoFisher 1:500; Mouse anti-protein A, Millipore Sigma 1:500; Mouse anti-Ty, ThermoFisher 1:1000) diluted in blocking solution (1% goat serum, 0.1% Triton X-100 in PBS) for 1 hour at room temperature or overnight at 4 °C. Cells were washed 3 times in PBS + 0.1% Tween-20 (PBST) and then incubated with secondary antibody (goat anti-rabbit Alexa 488, 594, or 647; ThermoFisher) diluted 1:1000 in blocking solution. Prior to mounting in Vectashield (VectorLabs), cells were washed and stained with 0.2 µg/mL DAPI in PBS for 5 minutes. The slides were imaged on a Leica SP5-II with a 100× objective. Z-series were obtained for each fluorescence channel and brightfield. For fluorescence images, maximum projections are shown. For indicated images, deconvolution analysis was performed on the fluorescent images using Huygens Essential deconvolution software (version 17.04.1p2 64b; SVI). Brightfield images are single confocal sections with a flatfield correction applied. All induced images were acquired with the same microscope settings as non-induced and parental controls.

### Western blotting

1E7 cells from log-phase cultures were centrifuged, washed in PBS and resuspended in Laemmli SDS-PAGE sample buffer at a concentration of 1×10^5^ cell equivalents/µL. Samples were boiled for 5 minutes at 95 °C and centrifuged briefly. Cell equivalents of 1.5×10^6^ cells per lane were separated by 10% SDS-PAGE with Precision Plus Protein Kaleidoscope (BioRad) as the standard. Fractionated proteins were transferred to a PVDF membrane and incubated in blocking buffer (PBS containing 5% milk and 0.2% Tween-20) for 1 hour at room temperature. Peroxidase anti-peroxidase soluble complex antibody (PAP reagent, Millipore Sigma 1:5000), rabbit anti-DLP antibody (1:1000), rabbit anti-aldolase (1:10000), mouse anti-Ty (1:1000), mouse anti-c-myc (1:1000), or mouse anti-alpha tubulin (KMX-1, Millipore Sigma, 1:1000) served as primary antibodies and were diluted in blocking buffer and added to blots prior to incubation overnight at 4 °C. Blots requiring a secondary antibody (all but the PAP reagent) were washed and probed with horseradish peroxidase-conjugated goat anti-mouse or anti-rabbit IgG in blocking buffer at 1:5000. After washing, blots were developed with chemiluminescent substrate (BioRad) and imaged on an Alpha Innotech or an Azure biosystems c600 imaging system using chemiluminescence settings.

### Transmission Electron Microscopy (TEM)

To visualize the ultrastructural effects of DLP depletion and overexpression, *T. brucei* PCF (strain 29-13) cells, including *Tb*DLP1/2_RNAi (±dox, 24h) and *Tb*DLP1::Ty^++^ (±dox, 48h) lines, were prepared for TEM at the University of Pennsylvania Electron Microscopy Resource Lab. Approximately 10^8^ log-phase cells were harvested by centrifugation and chemically fixed in 0.1 M sodium cacodylate buffer (pH 7.4) supplemented with 2.5% glutaraldehyde and 2.0% paraformaldehyde. Initial fixation was performed at room temperature for 2 hours, followed by an overnight incubation at 4 °C. Following primary fixation, samples were washed in buffer and post-fixed for 1 hour in 2.0% osmium tetroxide containing 1.5% K_3_Fe(CN)_6_. After rinsing with distilled water, samples underwent *en bloc* staining with 2% uranyl acetate. Dehydration was performed through a graduated ethanol series, after which samples were infiltrated and embedded in Embed-812 resin (Electron Microscopy Sciences). Ultrathin sections were secondarily stained with uranyl acetate and SATO lead. Imaging was conducted using a JEOL 1010 electron microscope equipped with a Hamamatsu digital camera and AMT Advantage NanoSprint500 software.

## Results

### Dynamin-like protein of *C. fasciculata* associates with glycosomal proteins

In other eukaryotes, dynamin-related proteins associate with membranes via adaptor proteins (19). To identify adaptor proteins in trypanosomatids, we used a tandem-affinity purification approach. *Crithidia fasciculata* can be grown to high densities making it a useful model for protein purification. We amplified the *Cf*DLP sequence from genomic DNA and cloned it into a plasmid that would append a C-terminal PTP tag (43) (Fig. 1A). We then introduced this construct into our parental *C. fasciculata* strain, where it is maintained as an episome. As a control, we created cell lines that expressed a PTP-tagged RFC3 (replication factor C) or RRP4 (RNase protein 4, part of the exosome complex), two proteins that had been purified previously using this approach (43). Using western blotting to detect the protein A portion of the tag, we observed tagged bands of the expected size for each PTP-tagged cell line (Fig. 1B). To confirm that *Cf*DLP::PTP had a similar localization as untagged *Cf*DLP, we performed immunofluorescence (IF) using a mouse antibody to detect the protein A tag and a custom rabbit antibody designed to detect the endogenous DLP from *T. brucei* (Fig. S1). We observed good co-localization between these two signals that was punctate with an area of enrichment near the flagellar pocket (Fig. S1). We then subjected *C. fasciculata* cells expressing either *Cf*RRP4::PTP or *Cf*DLP::PTP to the tandem affinity purification procedure. Silver stain of the final eluates showed a distinct banding pattern between the two, with a noticeable doublet around ∼90 kDa specifically in the *Cf*DLP::PTP eluate, near the expected migration for *Cf*DLP (∼77.3 kDa untagged, Fig. 1C). A western blot using the endogenous anti-*Tb*DLP antibody confirmed that the protein was reduced in size following the TEV protease step (which removes a portion of the PTP tag) and enriched in the final eluate (Fig. 1D).

**Fig 1.**
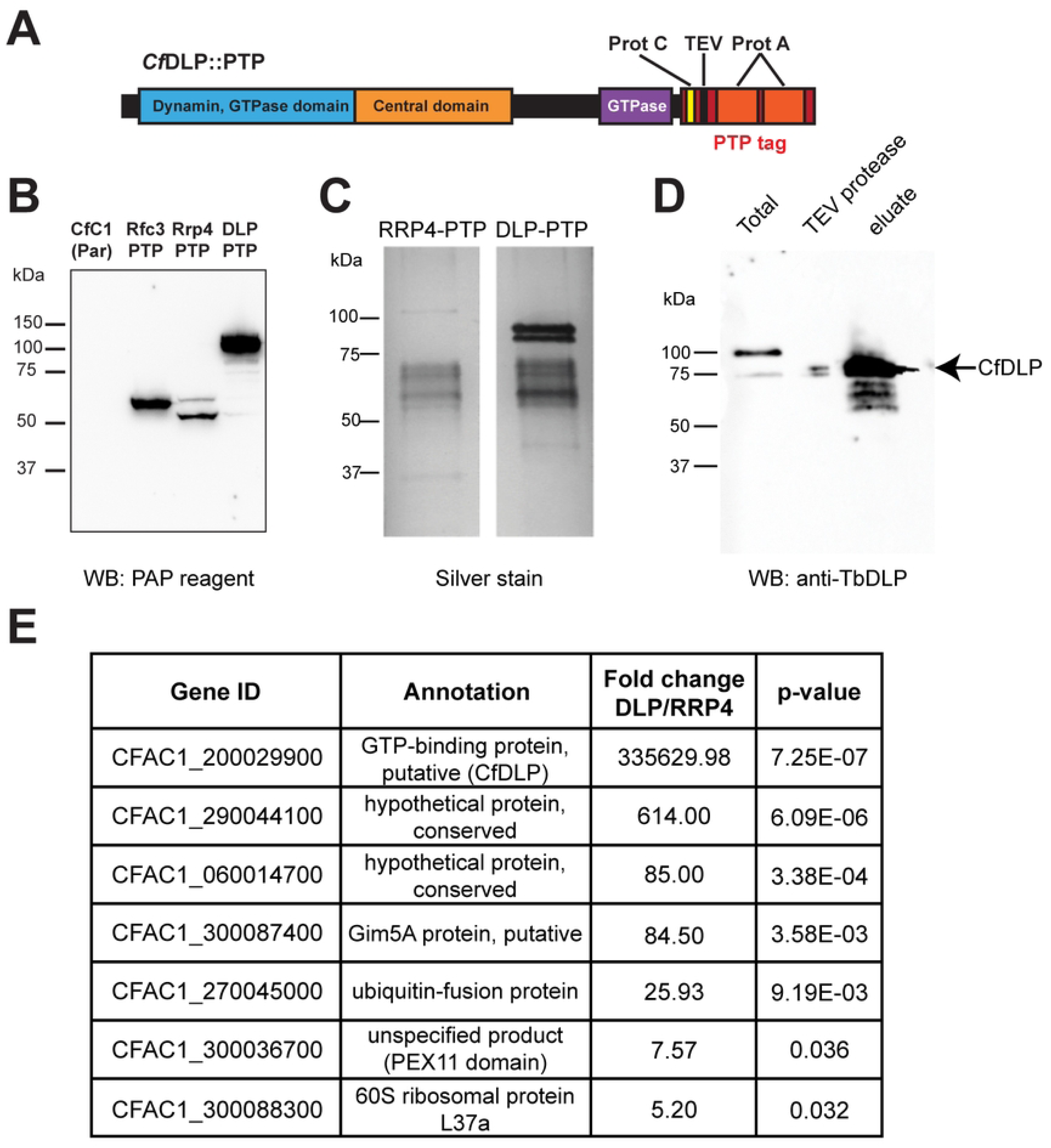
Purification of *Cf*DLP. **A**) Predicted domains of *Cf*DLP tagged with a C-terminal PTP-tag. **B**) Western blot of protein lysates prepared from parental cells (CfC1) and cells expressing various PTP-tagged proteins probed with the PAP reagent to detect the PTP tag. **C**) Silver stain of final eluates from purification of *Cf*RRP4::PTP and *Cf*DLP::PTP. **D**) Western blot of protein lysates obtained from the *Cf*DLP::PTP purification procedure probed with the antibody detecting endogenous DLP. In the total lysate lane, both PTP-tagged (upper band) and endogenous *Cf*DLP protein (lower band) can be seen. The middle lane shows a sample of the lysate following protein A-based purification and cleavage with TEV protease. The *Cf*DLP::PTP band has been reduced in size due to removal of a large portion of the tag. The final eluate lysate shows enrichment of *Cf*DLP. **E**) TriTrypDB accession numbers and annotations of the most highly enriched proteins detected in the *Cf*DLP eluate compared to the *Cf*RRP4 eluate.

We then subjected the *Cf*RRP4::PTP and *Cf*DLP::PTP eluates to quantitative label-free mass spectrometry. Proteins enriched in the *Cf*RRP4::PTP samples relative to the *Cf*DLP::PTP samples (p-value<0.05) included all nine expected subunits of the exosome complex (RRP4, RRP6, RRP40, RRP41A/ B, RRP45, EAP1, EAP2, EAP4, and CSL4) as well as two additional hypothetical conserved proteins (CFAC1_210027300 and CFAC1_180026700) and a putative mitochondrial phosphate transporter that had a relatively low enrichment (2.8 fold compared to >50-fold for the other hits) but a significant p-value (Table S2). Proteins enriched in *Cf*DLP::PTP samples relative to the *Cf*RRP4::PTP samples (p<0.05) included *Cf*DLP itself, predicted glycosomal proteins Gim5A and a PEX11-domain containing protein, two hypothetical conserved proteins (CFAC1_290044100 and CFAC1_060014700), a ubiquitin fusion protein, and a 60S ribosomal protein (Fig. 1E and Table S2). Other predicted glycosomal proteins were also enriched in the *Cf*DLP::PTP eluate but were above the p-value cutoff. These include a PEX11 ortholog (CFAC1_300036700), phosphoglycerate kinase (CFAC1_180006900), and another PEX11-domain containing protein (CFAC1_210006500, annotated as an unspecified product). Interestingly, about a third of the proteins identified are orthologous to known glycosomal proteins in *Leishmania* and *T. brucei* (66).

In *T. brucei*, *Tb*DLP1 is phosphorylated, which likely plays a role in its regulation (56,57). Although a protein alignment between *Cf*DLP, *Tb*DLP1 and *Tb*DLP2 show that the serine residues modified in TbDLP1 are not conserved in the *C. fasciculata* protein (Fig. 2A), we re-analyzed our mass spectrometry data to look for phosphorylated peptides (Table S3). We detected 14 phosphorylated proteins across all samples, including *Cf*DLP, with modifications detected at S260, S503, S592, and T594 in the amino acid sequence. We used AlphaFold3 to compare the predicted structures and locations of phosphorylation sites between *Tb*DLP1 and *Cf*DLP (Fig. 2B). In *Tb*DLP1, S537 and S540 are found in an unstructured, low confidence portion of the structure. In *Cf*DLP, the corresponding region is expanded and rich in Q, A, K, and D residues, with the predicted phosphorylation sites S503, S592, and T594 located at the boundaries of this region (Fig. 2).

**Fig 2.**
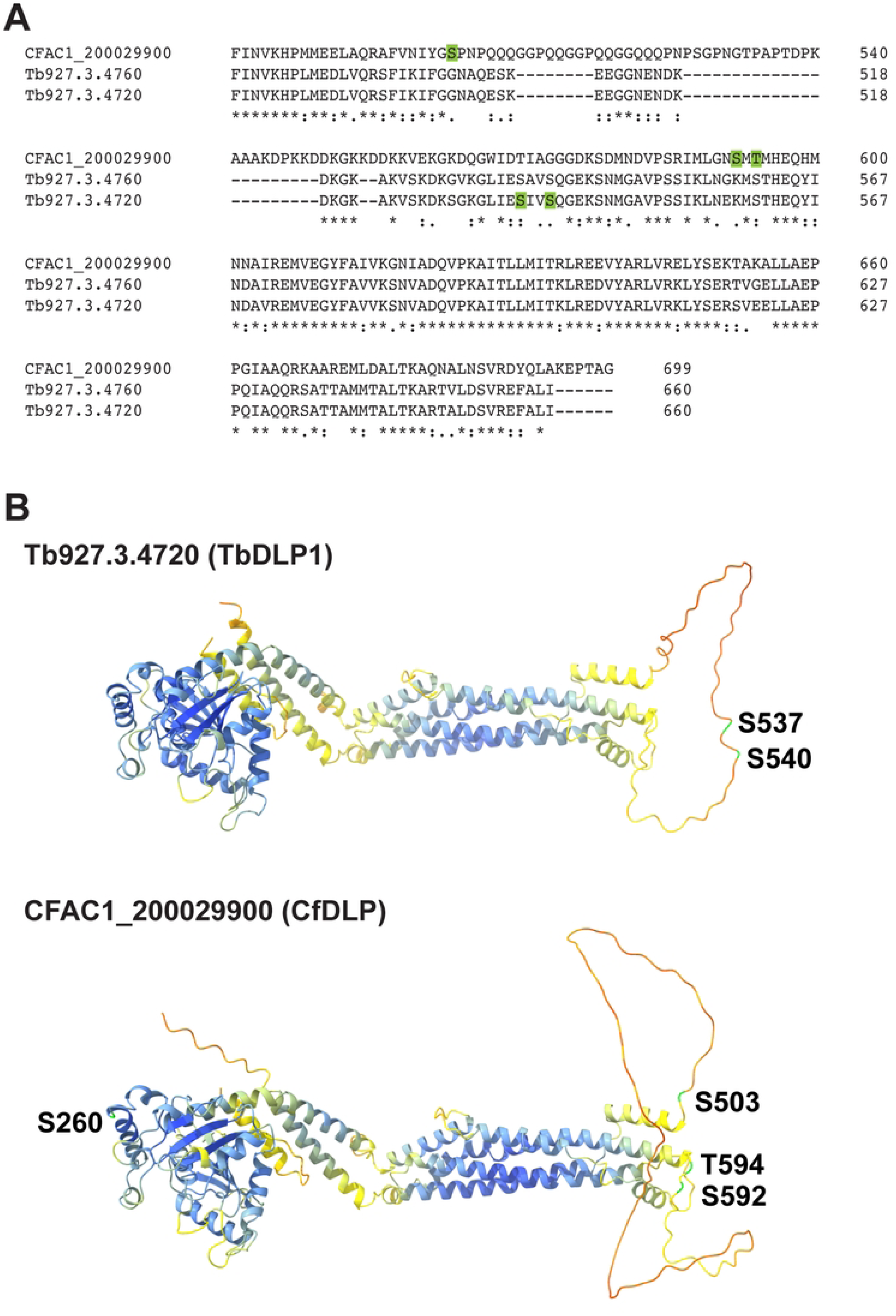
DLPs are phosphorylated. **A**) Alignment of the C-terminal region of *Cf*DLP (CFAC1_200029900), *Tb*DLP2 (Tb927.3.4760) and *Tb*DLP1 (Tb927.3.4720). Phosphorylated residues for *Tb*DLP1 (56,57) and *Cf*DLP (this study) are highlighted in green. **B**) AlphaFold structural predictions for *Cf*DLP and *Tb*DLP1 with phosphorylated residues labeled.

### DLP is found at constriction points on glycosomes

Since *Cf*DLP bound to glycosomal proteins in our purification experiments, we next sought to determine if *Cf*DLP co-localizes with glycosomes in the cell. To do this, we first established a glycosomal marker in *C. fasciculata* by tagging an episomal copy of *Cf*Gim5A with a C-terminal myc tag (*Cf*Gim5A::myc). Tagged *Cf*Gim5A::myc co-localizes with the IF signal for a glycosomal aldolase protein (Fig. S2). We then performed Ultrastructure Expansion Microscopy (U-ExM) on *Cf*Gim5A::myc-expressing cells and used IF to detect both the myc tag and endogenous *Cf*DLP (Fig. 3). The signal from tagged Gim5A again co-localized with glycosomes which were also visible by NHS ester staining. For endogenous *Cf*DLP, we observed a concentration of puncta around the flagellar pocket, as well as numerous cytoplasmic puncta (Fig. 3A-C). Some of these puncta were found on glycosomes, with stronger enrichment at points of constriction (Fig. 3D). In one case, *Cf*DLP appears to form spirals around a constricted glycosome (Fig. 3D, lower right).

**Fig 3.**
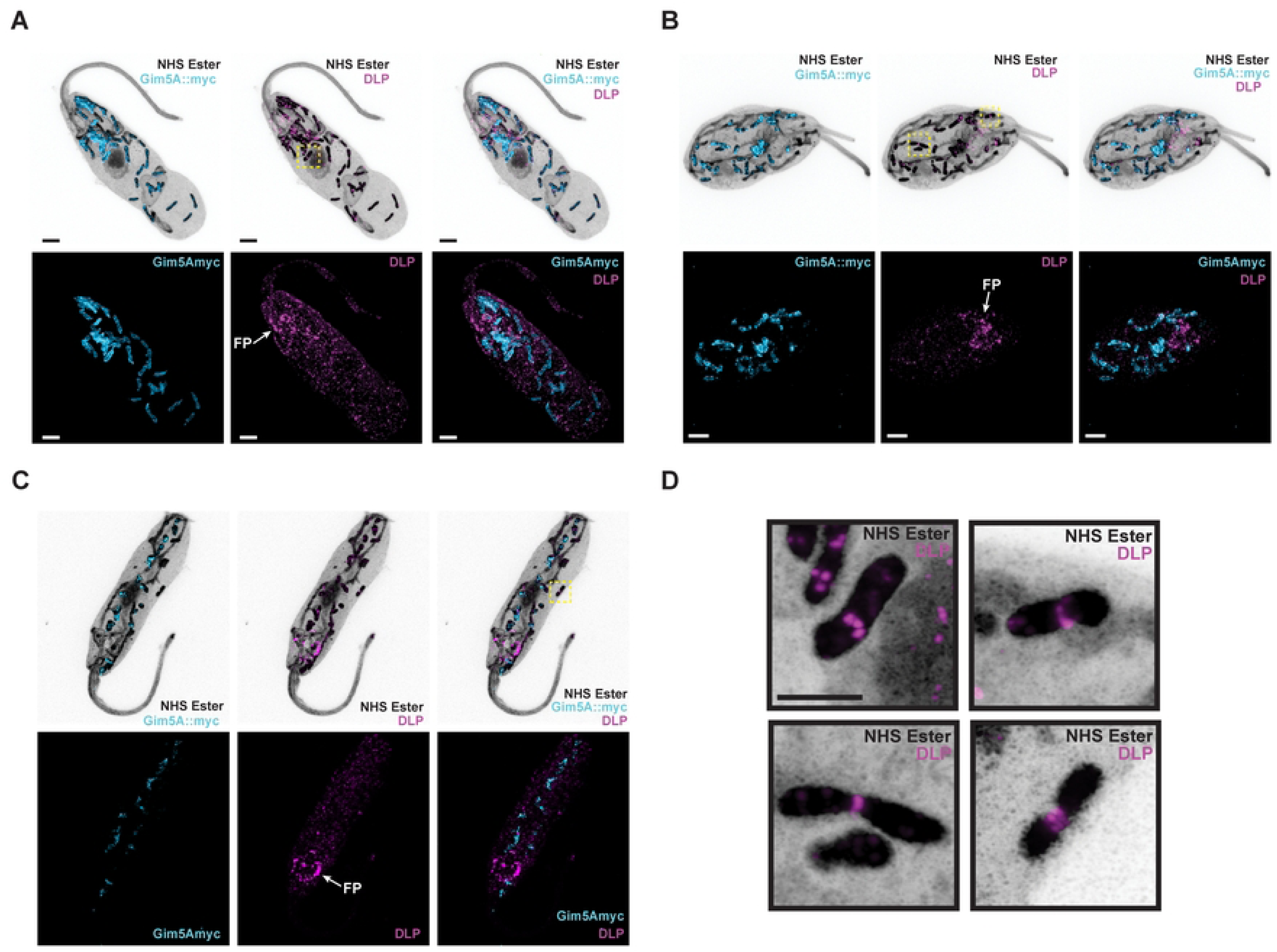
*Cf*DLP assembles around constricting glycosomes. **A-C**) CfC1 cells expressing *Cf*Gim5A::myc were expanded and stained with NHS ester (gray), α-myc antibodies (cyan), and α-DLP antibodies (magenta). FP, flagellar pocket. Scale bar 5 µm. **D**) Enlargements of the areas outlined with yellow dotted boxes. Scale bar 2.5 µm.

To see if DLP is found on glycosomes in other trypanosomatids, we performed a similar experiment in PCF *T. brucei*. Glycosomes were detected by introducing an ectopic construct driving expression of an enhanced yellow fluorescent protein (eYFP) fused to an N-terminal PTS2 glycosomal localization signal (Bauer et al., 2013). We again confirmed that the eYFP signal co-localizes with that of endogenous glycosomal aldolase (Fig. S2) before performing U-ExM on cells expressing PTS2::eYFP. The eYFP signal was detected with a mouse anti-GFP antibody, while the *Tb*DLP signal was detected using a rabbit antibody to the endogenous protein. In *T. brucei*, glycosomes were smaller and more numerous, with fewer obvious constrictions (Fig. 4A-B). *Tb*DLP signal was punctate, with brighter and larger puncta near the flagellar pocket. While some of the cytoplasmic *Tb*DLP puncta did co-localize with glycosomes (Fig. 4C-D), it was difficult to tell if this overlap was specific or correlated with any particular glycosomal morphology.

**Fig 4.**
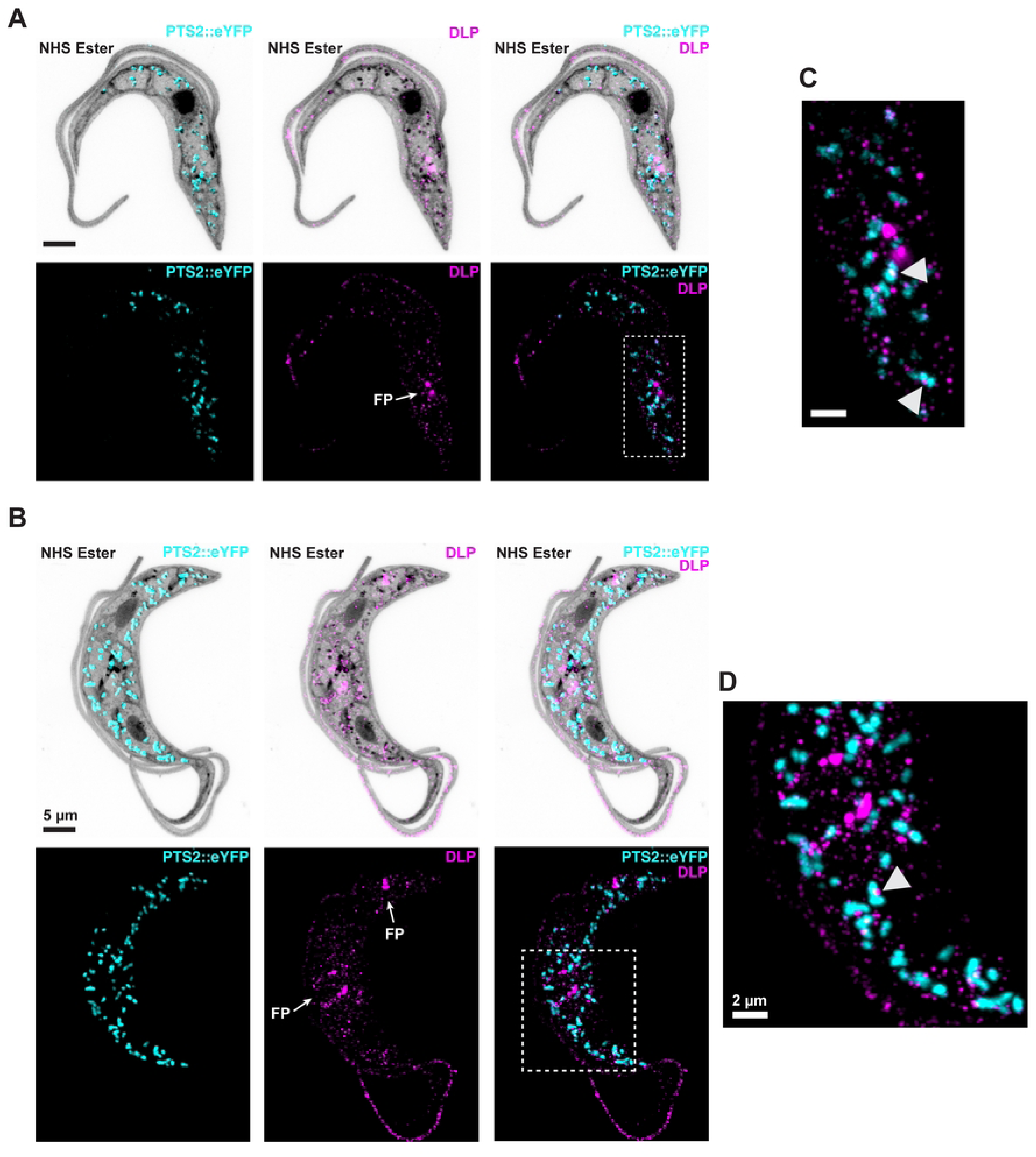
Localization of *Tb*DLP in *T. brucei*. **A-B**) PCF *T. brucei* cells expressing a construct with eYFP fused to a glycosomal targeting signal (PTS2::eYFP, cyan) were expanded and stained with NHS ester (gray) and α-DLP antibodies (magenta). FP, flagellar pocket. Scale bar 5 µm. **C-D**) Enlargements of areas outlined in gray dotted boxes. Arrowheads show points of co-localization between glycosomal and α-DLP signal. Scale bar 2 µm.

### Effects of *Tb*DLP levels on glycosome morphology

To see if depletion of DLP causes changes in glycosome morphology, we performed RNAi-mediated knockdown in PCF *T. brucei*. A construct driving doxycycline (dox)-inducible expression of a stem-loop double-stranded RNA construct targeting both *Tb*DLP1 and *Tb*DLP2 was previously shown to cause cell death, an endocytosis defect including a dramatically enlarged flagellar pocket, and a 2N2K cell cycle arrest due to a lack of mitochondrial fission and a resulting cytokinesis block (Chanez et al., 2006). Because *Tb*DLP1/2 knockdown causes dramatic changes to the overall morphology of the cell, we chose to examine cells after 24 hours of *Tb*DLP knockdown to avoid confounding phenotypes. In our cell line, we confirmed the growth phenotype and knockdown of the *Tb*DLP protein (Fig 5A-B). Antibody raised against *Tb*DLP1 detected a doublet by western blotting, with the lower band noticeably reduced in the presence of doxycycline. While both of these bands are within the range of the predicted molecular weights for *Tb*DLP1 and *Tb*DLP2, their similar size (73.3 kDa and 73.1 kDa respectively) makes their resolution into distinct bands unlikely. Levels of glycosomal aldolase remain unchanged during knockdown of *Tb*DLP.

**Fig 5.**
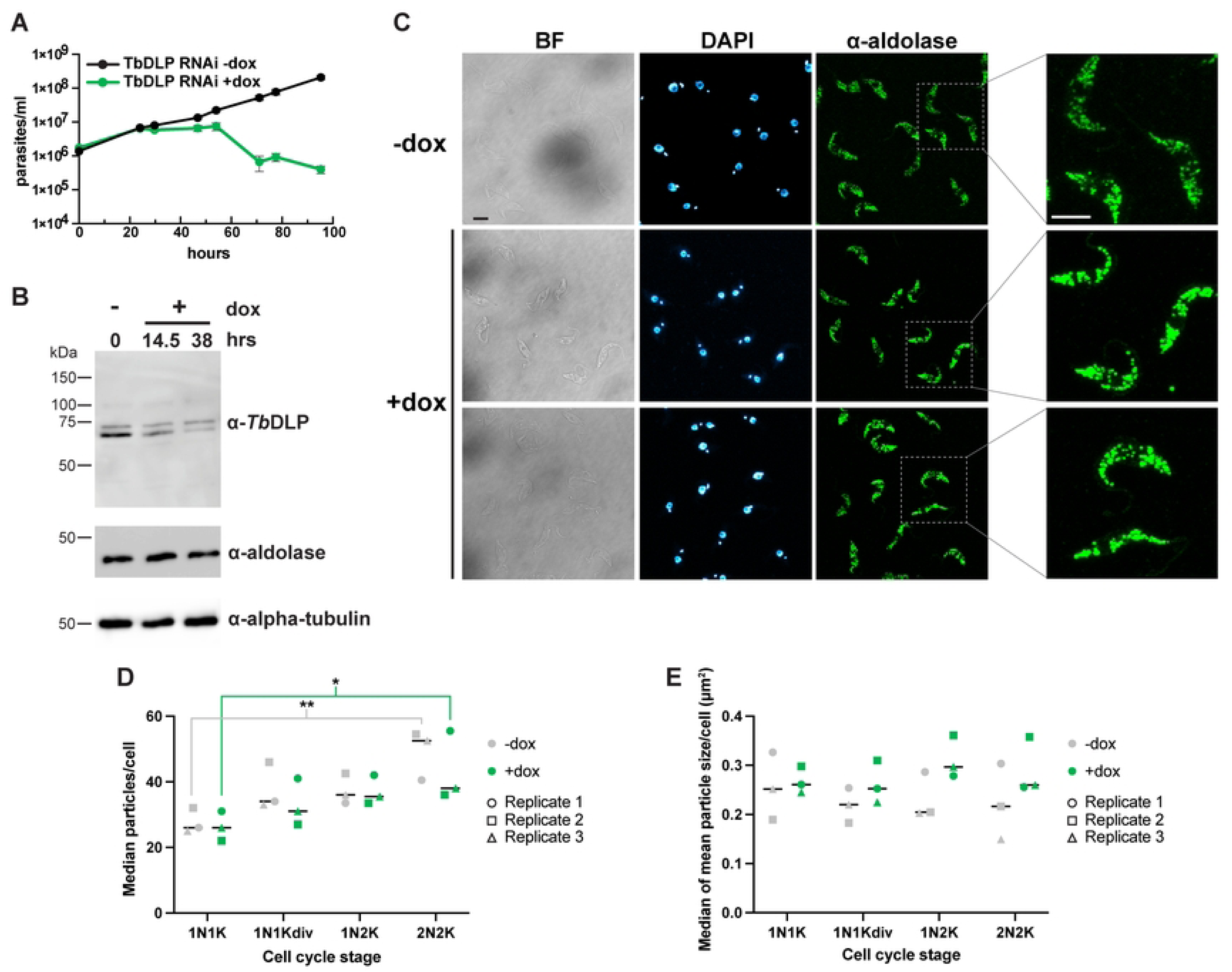
RNAi-mediated knockdown of *Tb*DLP impacts glycosomes. **A**) Growth curve of *T. brucei* PCF cells harboring the inducible *Tb*DLP1/2 RNAi construct grown in the absence (-dox, uninduced) or presence (+dox, induced) of doxycycline. Mean of three biological replicates. Error bars show standard deviation. **B**) Western blot of lysates prepared from *Tb*DLP RNAi cells grown in the absence (-) or presence (+) of doxycycline for the indicated times probed with antibodies to *Tb*DLP, aldolase, or alpha tubulin. Expected molecular weights are 73.3 kDa for *Tb*DLP1, 73.1 kDa for *Tb*DLP2, 41.1 kDa for aldolase, and 49.8 kDa for alpha-tubulin. **C**) Immunofluorescence microscopy with α-aldolase antibody comparing uninduced (-dox) cells and cells induced for RNAi of *Tb*DLP1/2 for 24 h. Right-hand column shows enlargement of the areas in the dotted boxes. BF, brightfield. Maximum projections are shown for aldolase and DAPI. Scale bars are 5 µm. **D**) ImageJ analysis of number of particles per cell detected with the anti-aldolase antibody as a function of cell cycle stage during knockdown of *Tb*DLP1/2. For each field, a threshold was applied to a maximum projection. ROIs were drawn around individual cells, which were scored and categorized for cell cycle stage based on the number of nuclei (N) and kinetoplasts (K) and whether the kinetoplast was dividing (K_div_). Each symbol represents the median number of particles per cell for that cell cycle stage in a biological replicate. N>200 cells per replicate. \*\**P* < 0.01, \**P* < 0.05, Two-way ANOVA, Tukey’s multiple comparisons test. **E**) As part of the same analysis that produced panel D, the mean particle size per cell was calculated. The median of these values is shown for each cell cycle stage and biological replicate. Two-way ANOVA revealed a significant global effect of *Tb*DLP RNAi on glycosome size (*P* = 0.03) although a Tukey’s multiple comparison test did not identify a specific cell cycle stage where this difference reached significance.

We then visualized glycosomes in cells undergoing *Tb*DLP knockdown by performing IF with anti-aldolase antibodies (Fig 5C). At 24 hours, we did not observe drastic changes to glycosomes, although in some cells glycosomes appeared slightly larger and brighter (Fig 5C). To measure changes in the distribution of aldolase signal following *Tb*DLP knockdown, we analyzed particle sizes in maximum projections of confocal z-stacks. Although not a direct measurement of glycosomes, since some three-dimensional information is lost, this approach should reveal if glycosomes fail to divide, resulting in fewer and larger aldolase particles. When we compared the median number of particles per cell across cell cycle stages, we observed no significant difference between uninduced (-dox) and induced (+dox) *Tb*DLP RNAi cells (Fig 5D). However, cell cycle stage did affect median particle number [Two-way ANOVA, *P* = 0.0014; F(3,16)=8.4] suggesting possible changes in organelle number or distribution as cells increase in size prior to division. For both-dox and + dox cells, the median number of aldolase particles/cell was lower in G1 phase cells (having 1 nucleus and 1 kinetoplast, or 1N1K) compared to G2 phase cells (those that have divided both the nucleus and kinetoplast, or 2N2K; two-way ANOVA with Tukey’s multiple comparisons test; *P* = 0.0055 for uninduced and *P* = 0.0311 for induced). We then calculated the mean particle size per cell and compared median values for uninduced and induced samples (Fig 5E). While there was a subtle but statistically significant global increase in particle size between uninduced and induced cells (Two-way ANOVA, *P* = 0.03), post-hoc multiple comparison tests did not identify a specific cell cycle stage where this difference reached significance.

To examine the morphology of *Tb*DLP RNAi cells in more detail, we performed transmission electron microscopy (TEM). We did not observe gross changes to the morphology of glycosomes, although they were difficult to identify unambiguously (Fig 6). We noted both smaller particles bound by a single membrane as well as larger particles that were usually more electron dense both in their lumen and their surrounding membrane. These larger particles stained more faintly in induced (+dox) *Tb*DLP RNAi cells. Although these cells were only induced for 24 hours, some already had enlarged flagellar pockets and mitochondrial abnormalities such as increased constrictions as previously reported (Benz et al., 2017).

**Fig 6.**
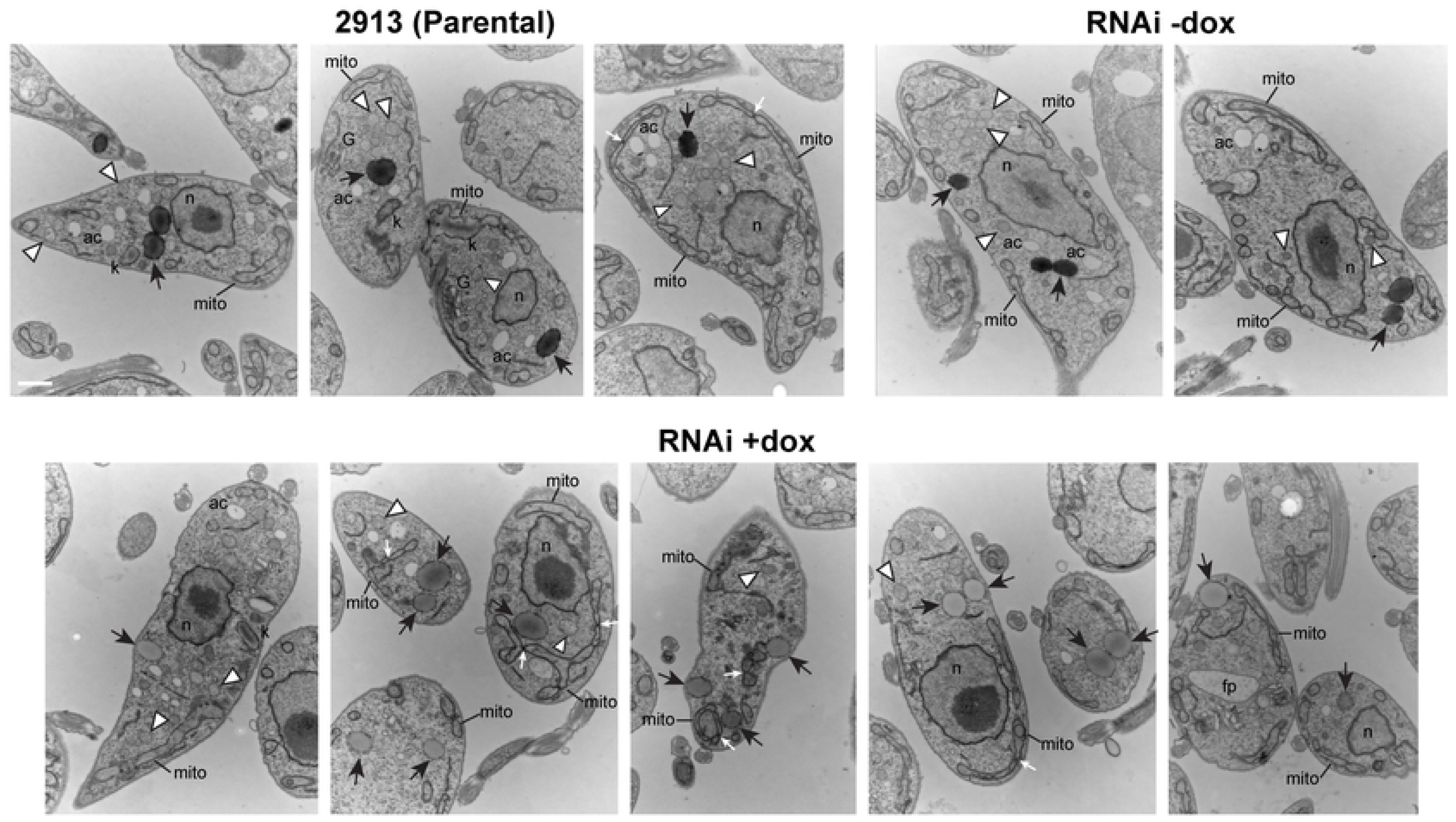
Ultrastructural changes during RNAi-mediated knockdown of *Tb*DLP1/2. Transmission electron micrographs of parental (29–13) cells, uninduced (-dox) *Tb*DLP1/2 RNAi cells and *Tb*DLP1/2 RNAi cells that had been induced (+dox) for 24 hours. N, nucleus; mito, mitochondrion; k, kinetoplast DNA; ac, acidocalcisome; G, Golgi; black arrows, large, electron-dense cytoplasmic vesicles; white arrowheads, small, heterogeneously-stained cytoplasmic vesicles. Both types of vesicles may represent glycosomes. Scale bar for all images is 800 nm.

Since analysis of *Tb*DLP RNAi cells is complicated by the pleiotropic nature of the knockdown phenotype, we asked whether overexpression of *Tb*DLP would alter glycosomal morphology. We cloned *Tb*DLP1 into a plasmid that adds a C-terminal Ty tag and allows for expression of this tagged *Tb*DLP1::Ty in the presence of dox. Overexpression of wild-type *Tb*DLP1::Ty did not noticeably impair growth (Fig 7A), although we did achieve robust expression as detected by both the anti-Ty and anti-endogenous DLP antibodies (Fig 7B-C). Probing western blots created with overexpression (*Tb*DLP1::Ty^++^) lysates with endogenous anti-*Tb*DLP antibodies again showed a doublet, but only the lower band appears overexpressed. The same blots probed with an antibody to the Ty epitope tag showed a single band in +dox lanes.

**Fig 7.**
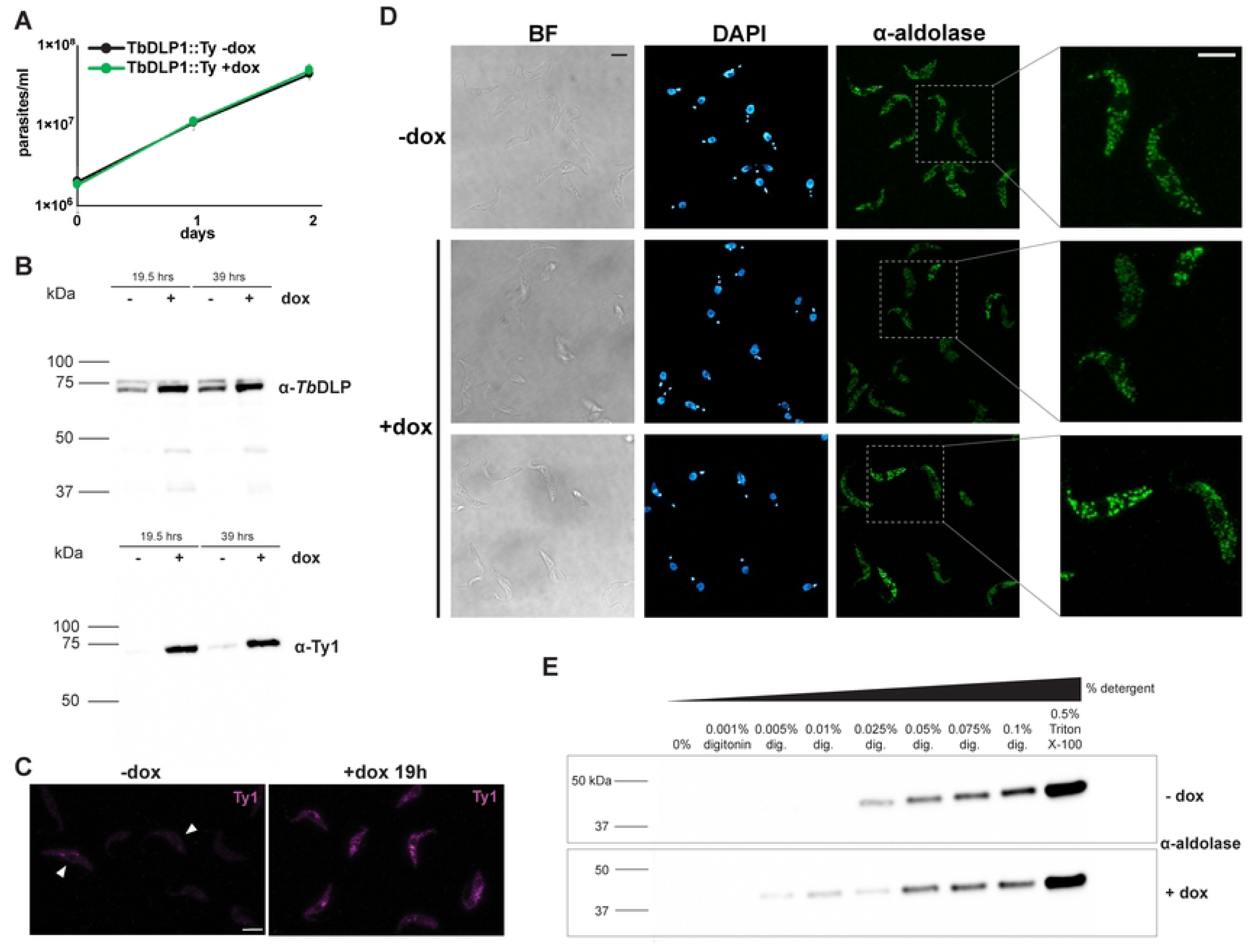
Overexpression of TbDLP1::Ty causes changes in glycosome permeability. A) Growth curve of *T. brucei* PCF cells harboring the inducible *Tb*DLP1::Ty overexpression construct grown in the absence (-dox, uninduced) or presence (+dox, induced) of doxycycline. Mean of three biological replicates. Error bars show standard deviation. **B**) Western blots confirm inducible overexpression of *Tb*DLP1::Ty detected with either an antibody raised again the endogenous *Tb*DLP (α-*Tb*DLP) or the C-terminal Ty epitope tag (α-Ty). **C**) Immunofluorescence microscopy with the α-Ty antibody in uninduced (-dox) and induced (+dox 19h) TbDLP::Ty^++^ cells. Maximum projections are shown. Scale bar is 5 µm. **D**) Immunofluorescence microscopy with the α-aldolase antibody comparing uninduced (-dox) and cells induced for overexpression of *Tb*DLP1::Ty for 48h. Right-hand column shows enlargement of the areas in the dotted boxes. BF, brightfield. Maximum projections are shown for aldolase and DAPI. Scale bars are 5 µm. **E**) Cell pellets from cells uninduced (-dox) or induced (+dox) for *Tb*DLP1::Ty for 50 h were solubilized with the indicated concentrations of detergent. Following centrifugation, supernatants were subjected to western blotting with α-aldolase antibodies. Additional replicates are shown in Fig S3.

To detect changes in glycosomes during *Tb*DLP1::Ty overexpression, we performed IF with the anti-aldolase antibody after 48 hours of dox induction (Fig 7D). We again did not observe dramatic changes in glycosomes, although in some cells the signal appeared more diffuse. To probe this further, we lysed cells with different concentrations of detergent to examine the solubility of a glycosomal marker (Fig 7E and S3). Overexpression of *Tb*DLP::Ty caused glycosomal aldolase to be released into the supernatant at lower detergent concentrations compared to lysis of -dox control cells.

We also used TEM to examine cells overexpressing *Tb*DLP1::Ty (Fig 8). While some cells appeared normal, others were vesiculated, with an increased number of smaller, membrane-bound compartments. Beyond this, we were unable to detect specific changes in glycosomes, the mitochondrion, or the flagellar pocket as a result of *Tb*DLP1 overexpression in PCF cells.

**Fig 8.**
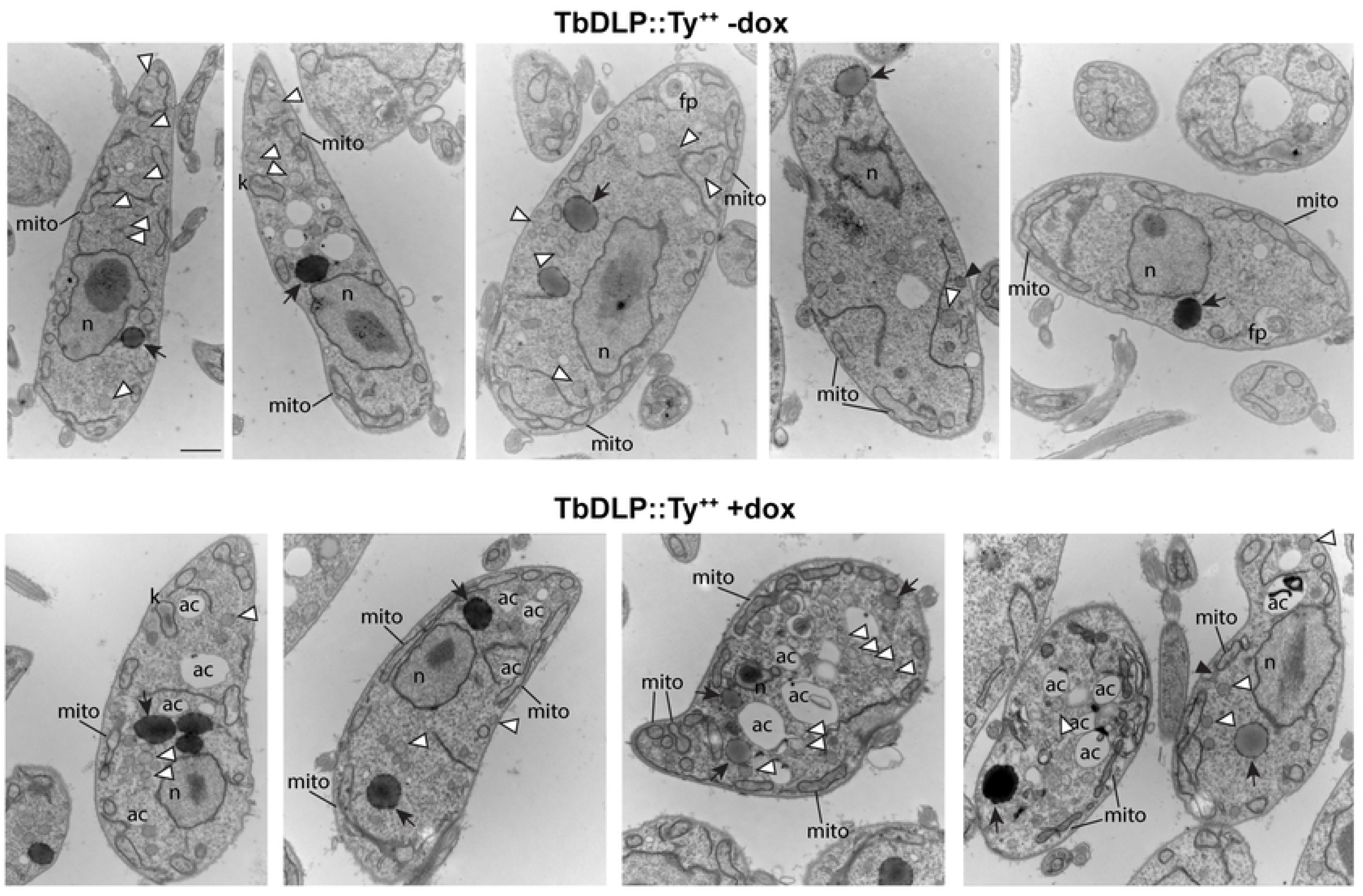
Ultrastructural changes during overexpression of *Tb*DLP1::Ty. Transmission electron micrographs of uninduced (-dox) *Tb*DLP1::Ty overexpression cells and *Tb*DLP1::Ty overexpression cells that had been induced (+dox) for 48 hours. N, nucleus; mito, mitochondrion; k, kinetoplast DNA; ac, acidocalcisome; G, Golgi; black arrows, large, electron-dense cytoplasmic vesicles; white arrowheads, small, heterogeneously-stained cytoplasmic vesicles. Both types of vesicles may represent glycosomes. Scale bar for all images is 800 nm.

## Discussion

Dynamin-like proteins in trypanosomatids are multifunctional proteins responsible for scission of endocytic vesicles and organelle division (**31–33**). These parasites are highly organized and undergo complex cell and developmental cycles, requiring precise regulation of DLP localization and function. In *Saccharomyces cerevisiae*, a dynamin-related protein called Vps1 is also required for both peroxisomal fission and endocytosis (67,68), suggesting that regulation of a single DLP at different cellular membranes may be a conserved feature of eukaryotic lineages.

Trypanosomatid DLPs lack pleckstrin homology domains and are likely recruited to membranes via specific adaptor proteins. In other organisms, adaptors can regulate DLP conformation (69) and/or function (70,71) or have their own membrane remodeling activities (72,73). While *Tb*DLP may interact with one or more adaptors at different cellular locations, there is also evidence that *Tb*DLP1 is phosphorylated and that this modification is more abundant in BSF parasites (56,57). Phosphorylation of serine 540 is significantly less abundant in *Tb*DYRK null mutants, suggesting that *Tb*DLP1 may be a substrate of this key differentiation kinase (74).

To add complexity, *T. brucei* has two, closely related *Tb*DLPs. *Tb*DLP1 is enriched in BSF cells and is sufficient to rescue the *Tb*DLP1/2 RNAi phenotype in this life cycle stage (31). In PCF, *Tb*DLP1 is cell-cycle regulated, peaking just prior to cell division (75). Although *Tb*DLP2 is more abundant in PCF cells, neither *Tb*DLP1 nor *Tb*DLP2 can fully reverse the effects of *Tb*DLP1/2 RNAi in PCF (31). How interacting proteins, post-translational modifications, and differential abundance of *Tb*DLP paralogs intersect to regulate dynamin activity on different cellular membranes is not yet understood.

*Crithidia fasciculata*, like other trypanosomatids in the *Leishmania* lineage, has only one predicted DLP. Perhaps the high endocytic rate of *T. brucei*, required for survival in its mammalian host, favored specialization of DLP function across two paralogous proteins. In this study, we have shown for the first time that *Cf*DLP, similar to *Tb*DLP1 and 2 (31), localizes in cytoplasmic puncta with a concentration around the flagellar pocket, the site of endocytosis. We have also shown that *Cf*DLP is phosphorylated in a region analogous to that of *Tb*DLP1 according to structural predictions. This suggests that important aspects of DLP function and regulation are likely conserved across trypanosomatids.

Probing western blots containing *T. brucei* lysates with our anti-*Tb*DLP antibody revealed two bands running as a near doublet in the ∼75 kDa range, close to the predicted size of DLP. These could represent *Tb*DLP1 and 2; however, we consider this unlikely as the two proteins differ by only 19 amino acids (a predicted size difference of 173 Da). Our purification of *Cf*DLP also produced a doublet in the final eluate, which could be due to self-association of the exogenous PTP-tagged *Cf*DLP with native *Cf*DLP. Self-assembly is a conserved feature of dynamins (76) and has been demonstrated for DLP in *L. donovani* (36). In our RNAi experiments in *T. brucei*, only the lower band is significantly knocked down, although the construct should target both *Tb*DLP1 and 2. We designed our overexpression construct to express only tagged *Tb*DLP1, which resulted in increased intensity of the lower band with the upper band again unaffected. We surmise that the upper band is either non-specific or represents a modified version of *Tb*DLP1. However, this modified version would have to be stable enough to persist following transcript knockdown by RNAi and fail to increase with overexpression. Interestingly, a previous study also appears to show a doublet when either PTP-tagged *Tb*DLP1 or *Tb*DLP2 is over-expressed (31). DLP from parental *C. fasciculata* does not run as a doublet.

Using co-immunoprecipitation, we sought to identify *Cf*DLP-interacting proteins, including adaptors, that could recruit this protein to the flagellar pocket and/or mitochondrial membrane. Recently, a similar approach successfully identified an anchor protein for Dyn2 in *Plasmodium falciparum* named *Pf*Anchor (27). Dyn2 is required for both mitochondrial fission as well as division of the plastid-like organelle called the apicoplast (29). Immunoprecipitation of Dyn2 revealed both *Pf*Anchor and a possible dynamin adaptor for the mitochondrion, an AAA-ATPase (27). In our experiments, we found the putative *Cf*Pex11 in addition to the PEX11-related protein Gim5A (77). Dynamin-related proteins in mammals and yeast mediate peroxisomal fission and can use Pex11 as an adaptor (78,79). In fact, dynamins can interact with the same adaptors on both mitochondrial and peroxisomal membranes, including Mff, Fis1 and Pex11p (80,81). As glycosomes are modified peroxisomes, it is therefore possible that Pex11 and/or Gim5A serve as dynamin adaptors on one or more organelles in trypanosomatids. Supporting a role for DLP in glycosomal fission, *Tb*DLP1/2 were discovered in a glycosomal proteome, although these samples contained some mitochondria (82). A recent preprint, published while this manuscript was in preparation, argues for co-fractionation of *Tb*DLP and the putative adaptor *Tb*Fis1 with glycosomes (83). There is no obvious ortholog for Fis1 in *C. fasciculata*, although it may be sufficiently diverged to prevent sequence-based identification. No proteins with a similar domain organization were detected in our IP samples.

Expansion microscopy, particularly in *C. fasciculata* which has larger and more distinct glycosomes, appears to show assembly of DLP around points of constriction. Future studies are needed to understand whether the GTPase activity of DLP is sufficient to produce these constrictions or whether additional proteins are required. In *T. brucei*, it has been proposed that glycosomes divide through fission of existing organelles or form *de novo* through vesicles that bud from the ER and fuse with other vesicles, resulting in glycosomal protein import and maturation (84). Whether DLP is also required for *de novo* glycosome formation is not known. In mammalian cells, vesicles derived from the mitochondrion also contribute to *de novo* peroxisome biogenesis (85).

Overall, we observed that the number of aldolase particles per cell increases during cell cycle progression, implying that glycosomal biogenesis may occur during cell growth to ensure even distribution during cell division. However, our experiments with *Tb*DLP1/2 RNAi cells did not show dramatic effects on glycosomes during protein knockdown. This may be due to time point selection, as flagellar pocket swelling and cell rounding precludes analysis of cellular structure past 24 hours of RNAi induction when glycosomal phenotypes might be observed. In addition, growth arrest may mask defects in glycosomal fission. Interestingly, we did observe a subtle trend towards larger glycosomes in *Tb*DLP1/2 knockdown cells at 24 hours which was not limited to a specific cell cycle stage.

Our TEM studies revealed phenotypes previously reported in *Tb*DLP RNAi cells, including mitochondrial constrictions (31,33). We note different categories of cytoplasmic vesicles, including ones that were relatively small and heterogeneously stained and larger, more uniformly-stained vesicles with an electron dense membrane. Based on the number of particles observed via immunofluorescence, both of these may represent sub-populations of glycosomes (12,13). In *Tb*DLP1/2 RNAi cells, we noted a change in the larger particles, as they appeared more electron lucent. As cells with reduced DLP have lower rates of endocytosis (32), alterations in membrane trafficking could delay glycosome maturation.

In contrast to *Tb*DLP1/2 knockdown, overexpression of *Tb*DLP1::Ty in PCF cells did not disrupt growth, consistent with a previous study in which wild-type *Tb*DLP was overexpressed (32). By IF, glycosomes appeared largely normal, although the signal in some cells was faint and less distinct. This is consistent with a change in aldolase solubility in differential fractionation experiments. By TEM, some *Tb*DLP1::Ty overexpression cells had increased numbers of intracellular vesicles. In these cells, increased endocytic traffic, excess vesicle budding, or increased fission may disrupt intracellular membranes, leading to changes in glycosomal membrane composition and increased sensitivity to detergent permeabilization. Overexpression of *Tb*DLP1 may fail to produce specific organelle alterations because its adaptors or other interacting partners are present in insufficient amounts to drive an increase in *Tb*DLP1 function.

The finding of a third putative function for the trypanosomatid DLP points to this protein as a master regulator of cellular organization in these important parasites. Intriguingly, *Tb*DLP1/2 has also been identified as a potential interaction partner of *Tb*PH1, a pleckstrin homology protein found on the microtubule quartet structure (86), and of TOEFAZ1, part of the cytokinesis machinery (87). Future studies across species will reveal how DLP accomplishes diverse membrane remodeling reactions to enable the survival and development of trypanosomatids across varied host niches.

## Acknowledgments

The authors thank Stuart A. MacNeill for providing the pNUS-PTPcH-CfRfc3, pNUS-PTPcH-CfRrp4, and unmodified pNUS-PTPcH constructs as well as advice and protocols. We also thank Emmanuel Tetaud for creating and providing the pNUS plasmids. We thank Meredith Morris for providing the aldolase antibody and PTS2eYFP construct and Andre Schneider for providing the *Tb*DLP1/2 inducible RNAi construct. Transmission electron microscopy was performed by Biao Zuo at the Electron Microscopy Resource Lab (EMRL) at the Perelman School of Medicine, University of Pennsylvania.

---

**Figure S1.** Expression and localization of *Cf*DLP. **A**) Detection of *Cf*DLP using an antibody to endogenous protein. Protein lysates from the *C. fasciculata* parental cell line (CfC1) and the parental cell line to the *Tb*DLP1/2 RNAi cells [TbPCFmitoGFP_E10, (47)] were run on an SDS-PAGE gel, transferred to PVDF membrane, and probed with a rabbit antibody raised against a portion of *Tb*DLP2. Predicted size of *Cf*DLP is 77.3 kDa. Predicted size of *Tb*DLP1 is 73.3 kDa. Predicted size of *Tb*DLP2 is 73.1 kDa. **B**) Immunofluorescence analysis of parental CfC1 cells and cells expressing the *Cf*DLP::PTP construct using antibodies against protein A (α-ProtA) to detect the tag and against DLP (α-DLP). DNA is stained with DAPI. BF, brightfield. Maximum projections are shown for fluorescent signals. Scale bar is 5 µm.

**Figure S2. Validation of glycosomal markers. A**) Immunofluorescence analysis of *C. fasciculata* cells expressing *Cf*Gim5A::myc stained with α-aldolase (magenta) and α-myc (green). DNA is stained with DAPI. Fluorescence images in the second row were subjected to Huygen’s deconvolution. Scale bar is 5 µm. **B**) Immunofluorescence analysis of *T. brucei* PCF cells expressing PTS2eYFPaldo (green) stained with α-aldolase (magenta). DNA was stained with DAPI. Scale bar is 5 µm.

**Figure S3. Overexpression of *Tb*DLP1::Ty alters the solubility of glycosomal aldolase. A**) Western blot of lysates obtained from *Tb*DLP1::Ty^++^ cells grown in the absence (-dox) or presence (+dox) for the indicated times. Blots were probed with antibodies detecting *Tb*DLP, aldolase, and alpha tubulin. **B**) Additional biological replicates of the experiment shown in Fig 7E. TbDLP1::Ty overexpression cells grown in the absence (-dox) or presence (+dox) of doxycycline were pelleted and resuspended in the indicated concentrations of detergent. Following centrifugation, supernatants were subjected to western blotting with an antibody raised against glycosomal aldolase. **C**) Quantitation of all three fractionation replicates using ImageJ. The signal in each lane was calculated as a percentage of the total signal in the digitonin-solubilized lanes. The Triton-X100 lane was not included in quantitation as its signal was outside the linear range.

## Notes

### Competing Interest Statement

The authors have declared no competing interest.

## References

1. World Health Organization. Global report on neglected tropical diseases 2025. 2025.

2. World Organisation for Animal Health (2025). The State of the World’s Animal Health 2025 [Internet]. Paris: WOAH (World Organisation for Animal Health); 2025 [cited 2026 Mar 4]. p. 120pp. Report No. Available from: https://www.woah.org/en/the-state-of-the-worlds-animal-health/doi:10.20506/woah.3586

3. Wheeler RJ, Gluenz E, Gull K. The Limits on Trypanosomatid Morphological Diversity. Li Z, editor. PLoS ONE. 2013 Nov 19;8(11):e79581. doi:10.1371/journal.pone.0079581

4. Wheeler RJ, Gull K, Sunter JD. Coordination of the Cell Cycle in Trypanosomes. Annu Rev Microbiol. 2019 Sep 8;73(1):133–54. doi:10.1146/annurev-micro-020518-115617

5. He CY, Ho HH, Malsam J, Chalouni C, West CM, Ullu E, et al. Golgi duplication in Trypanosoma brucei. J Cell Biol. 2004 May 10;165(3):313–21. doi:10.1083/jcb.200311076

6. Priest JW, Hajduk SL. Developmental regulation of mitochondrial biogenesis inTrypanosoma brucei. J Bioenerg Biomembr. 1994 Apr 1;26(2):179–91. doi:10.1007/BF00763067

7. Herman M, Pérez-Morga D, Schtickzelle N, Michels PAM. Turnover of glycosomes during life-cycle differentiation of Trypanosoma brucei. Autophagy. 2008 Apr 1;4(3):294–308. doi:10.4161/auto.5443 PubMed PMID: 18365344.

8. Zíková A. Mitochondrial adaptations throughout the Trypanosoma brucei life cycle. Journal of Eukaryotic Microbiology. 2022;69(6):e12911. doi:10.1111/jeu.12911

9. Allmann S, Bringaud F. Glycosomes: A comprehensive view of their metabolic roles in *T. brucei*. The International Journal of Biochemistry & Cell Biology. 2017 Apr 1;85:85–90. doi:10.1016/j.biocel.2017.01.015

10. Szöőr B, Simon DV, Rojas F, Young J, Robinson DR, Krüger T, et al. Positional Dynamics and Glycosomal Recruitment of Developmental Regulators during Trypanosome Differentiation. mBio. 2019 Jul 9;10(4): doi:10.1128/mbio.00875-19

11. Szöör B, Haanstra JR, Gualdrón-López M, Michels PA. Evolution, dynamics and specialized functions of glycosomes in metabolism and development of trypanosomatids. Current Opinion in Microbiology. 2014 Dec;22:79–87. doi:10.1016/j.mib.2014.09.006

12. Anderson H, Powell RR, Morris MT. Ultrastructural expansion microscopy reveals unexpected levels of glycosome heterogeneity in African trypanosomes. Journal of Microscopy. 2026;301(2):222–40. doi:10.1111/jmi.70019

13. Crowe LP, Morris MT. Glycosome heterogeneity in kinetoplastids. Biochem Soc Trans. 2021 Feb 26;49(1):29–39. doi:10.1042/BST20190517 PubMed PMID: 33439256; PubMed Central PMCID: PMC7925000.

14. Halliday C, de Castro-Neto A, Alcantara CL, Cunha-e-Silva NL, Vaughan S, Sunter JD. Trypanosomatid Flagellar Pocket from Structure to Function. Trends in Parasitology. 2021 Apr 1;37(4):317–29. doi:10.1016/j.pt.2020.11.005

15. da Cunha-e-Silva NL, de Alcantara C L, Pereira MG, Souza WD. Three Hungry Tryps: the efficient endocytic pathway of pathogenic trypanosomatids. Trends in Parasitology. 2025 Jul 1;41(7):560–71. doi:10.1016/j.pt.2025.05.005 PubMed PMID: 40483243.

16. Link F, Borges AR, Jones NG, Engstler M. To the Surface and Back: Exo- and Endocytic Pathways in *Trypanosoma brucei*. Front Cell Dev Biol. 2021 Aug 6;9:720521. doi:10.3389/fcell.2021.720521

17. Morgan GW, Allen CL, Jeffries TR, Hollinshead M, Field MC. Developmental and morphological regulation of clathrin-mediated endocytosis in Trypanosoma brucei. J Cell Sci. 2001 Jul 15;114(14):2605–15. doi:10.1242/jcs.114.14.2605

18. Ramachandran R, Schmid SL. The dynamin superfamily. Current Biology. 2018 Apr;28(8):R411–6. doi:10.1016/j.cub.2017.12.013

19. Bui HT, Shaw JM. Dynamin Assembly Strategies and Adaptor Proteins in Mitochondrial Fission. Current Biology. 2013 Oct;23(19):R891–9. doi:10.1016/j.cub.2013.08.040

20. Breinich MS, Ferguson DJP, Foth BJ, van Dooren GG, Lebrun M, Quon DV, et al. A dynamin is required for the biogenesis of secretory organelles in *Toxoplasma gondii*. Curr Biol. 2009 Feb 24;19(4):277–86. doi:10.1016/j.cub.2009.01.039 PubMed PMID: 19217293; PubMed Central PMCID: PMC3941470.

21. Melatti C, Pieperhoff M, Lemgruber L, Pohl E, Sheiner L, Meissner M. A unique dynamin-related protein is essential for mitochondrial fission in Toxoplasma gondii. PLOS Pathogens. 2019 Apr 4;15(4):e1007512. doi:10.1371/journal.ppat.1007512

22. Heredero-Bermejo I, Varberg JM, Charvat R, Jacobs K, Garbuz T, Sullivan Jr. WJ, et al. TgDrpC, an atypical dynamin-related protein in Toxoplasma gondii, is associated with vesicular transport factors and parasite division. Molecular Microbiology. 2019;111(1):46–64. doi:10.1111/mmi.14138

23. Koreny L, Mercado-Saavedra BN, Klinger CM, Barylyuk K, Butterworth S, Hirst J, et al. Stable endocytic structures navigate the complex pellicle of apicomplexan parasites. Nat Commun. 2023 Apr 15;14(1):2167. doi:10.1038/s41467-023-37431-x

24. van Dooren GG, Reiff SB, Tomova C, Meissner M, Humbel BM, Striepen B. A Novel Dynamin-Related Protein Has Been Recruited for Apicoplast Fission in *Toxoplasma gondii*. Current Biology. 2009 Feb 24;19(4):267–76. doi:10.1016/j.cub.2008.12.048

25. Verhoef JMJ, Meissner M, Kooij TWA. Organelle Dynamics in Apicomplexan Parasites. mBio. 2021 Aug 31;12(4):e0140921. doi:10.1128/mBio.01409-21 PubMed PMID: 34425697; PubMed Central PMCID: PMC8406264.

26. Li H, Han Z, Lu Y, Lin Y, Zhang L, Wu Y, et al. Isolation and functional characterization of a dynamin-like gene from *Plasmodium falciparum*. Biochemical and Biophysical Research Communications. 2004 Jul 30;320(3):664–71. doi:10.1016/j.bbrc.2004.06.010

27. Blauwkamp JA, Rajaram K, Staggers SR, Harrigan O, Doud EH, Xu W, et al. An essential adaptor for apicoplast fission and inheritance in malaria parasites. Nat Commun. 2025 Nov 27;16(1):11325. doi:10.1038/s41467-025-66393-5

28. Charneau S, Dourado Bastos IM, Mouray E, Ribeiro BM, Santana JM, Grellier P, et al. Characterization of PfDYN2, a dynamin-like protein of *Plasmodium falciparum* expressed in schizonts. Microbes and Infection. 2007 Jun 1;9(7):797–805. doi:10.1016/j.micinf.2007.02.020

29. Morano AA, Xu W, Navarro FM, Shadija N, Dvorin JD, Ke H. The dynamin-related protein PfDyn2 is essential for both apicoplast and mitochondrial fission in *Plasmodium falciparum*. mBio. 2024 Nov 29;16(1):e03036–24. doi:10.1128/mbio.03036-24

30. Thakur V, Islam MdM, Singh S, Rathore S, Muneer A, Dutta G, et al. A dynamin-like protein in *Plasmodium falciparum* plays an essential role in parasite growth, mitochondrial development and homeostasis during asexual blood stages. Biochimica et Biophysica Acta (BBA) - Molecular Cell Research. 2025 Jun 1;1872(5):119940. doi:10.1016/j.bbamcr.2025.119940

31. Benz C, Stříbrná E, Hashimi H, Lukeš J. Dynamin-like proteins in Trypanosoma brucei: A division of labour between two paralogs? Mata J, editor. PLoS ONE. 2017;12(5):e0177200–19. doi:10.1371/journal.pone.0177200

32. Chanez AL, Hehl AB, Engstler M, Schneider A. Ablation of the single dynamin of *T. brucei* blocks mitochondrial fission and endocytosis and leads to a precise cytokinesis arrest. Journal of Cell Science. 2006;119(14):2968–74. doi:10.1242/jcs.03023

33. Morgan GW, Goulding D, Field MC. The Single Dynamin-like Protein of Trypanosoma brucei Regulates Mitochondrial Division and Is Not Required for Endocytosis. Journal of Biological Chemistry. 2004;279(11):10692–701. doi:10.1074/jbc.m312178200

34. Morel CA, Asencio C, Moreira D, Blancard C, Salin B, Gontier E, et al. A new member of the dynamin superfamily modulates mitochondrial membrane branching in *Trypanosoma brucei*. Current Biology. 2025 Mar 24;35(6):1337–1352.e5. doi:10.1016/j.cub.2025.02.033

35. Vanwalleghem G, Fontaine F, Lecordier L, Tebabi P, Klewe K, Nolan DP, et al. Coupling of lysosomal and mitochondrial membrane permeabilization in trypanolysis by APOL1. Nature Communications. 2015;6:8078. doi:10.1038/ncomms9078

36. Wuyts E, Sundaramoorthy R, Tulloch L, Monsieurs P, Eadsforth TC, Pintelon I, et al. Biophysical analysis of an oligomerization-attenuated variant of the *Leishmania donovani* dynamin-1-like protein. Molecular and Biochemical Parasitology. 2025 Sep 1;263:111691. doi:10.1016/j.molbiopara.2025.111691

37. Hefnawy A, Negreira G, Jara M, Cotton JA, Maes I, D’Haenens E, et al. Genomic and Phenotypic Characterization of Experimentally Selected Resistant *Leishmania donovani* Reveals a Role for Dynamin-1-Like Protein in the Mechanism of Resistance to a Novel Antileishmanial Compound. Soldati-Favre D, editor. mBio. 2022 Feb 22;13(1):e03264–21. doi:10.1128/mbio.03264-21

38. DiMaio GRJJCMFMMLPJ. The single mitochondrion of the kinetoplastid parasite Crithidia fasciculata is a dynamic network. PLoS ONE. 2018;13(12):1–21. doi:10.1371/journal.pone.0202711

39. Warren WC, Akopyants NS, Dobson DE, Hertz-Fowler C, Lye LF, Myler PJ, et al. Genome Assemblies across the Diverse Evolutionary Spectrum of *Leishmania* Protozoan Parasites. Microbiology Resource Announcements. 2021 Sep 2;10(35):10.1128/mra.00545-21. doi:10.1128/mra.00545-21

40. Bieber BV, St Clair FA, Quatse J, Denecke S, Povelones M, Povelones ML. Culture and Genetic Modification of the Monoxenous Trypanosomatids Crithidia fasciculata and Crithidia bombi. Methods Mol Biol. 2026;3013:73–87. doi:10.1007/978-1-0716-5142-1_5 PubMed PMID: 41627731.

41. Wirtz E, Hoek M, Cross GA. Regulated processive transcription of chromatin by T7 RNA polymerase in Trypanosoma brucei. Nucleic Acids Res. 1998 Oct 15;26(20):4626–34. doi:10.1093/nar/26.20.4626 PubMed PMID: 9753730; PubMed Central PMCID: PMC147901.

42. Wirtz E, Clayton C. Inducible gene expression in trypanosomes mediated by a prokaryotic repressor. Science. 1995 May 26;268(5214):1179–83. doi:10.1126/science.7761835 PubMed PMID: 7761835.

43. Kipandula W, Smith TK, MacNeill SA. Tandem affinity purification of exosome and replication factor C complexes from the non-human infectious kinetoplastid parasite *Crithidia fasciculata*. Molecular and Biochemical Parasitology. 2017 Oct;217:19–22. doi:10.1016/j.molbiopara.2017.08.004

44. Tetaud E, Lecuix I, Sheldrake T, Baltz T, Fairlamb AH. A new expression vector for *Crithidia fasciculata* and *Leishmania*. Molecular and Biochemical Parasitology. 2002 Apr;120(2):195–204. doi:10.1016/S0166-6851(02)00002-6

45. Bauer S, Morris JC, Morris MT. Environmentally Regulated Glycosome Protein Composition in the African Trypanosome. Eukaryotic Cell. 2013 Jul 25;12(8):1072–9. doi:10.1128/ec.00086-13

46. Burkard GS, Jutzi P, Roditi I. Genome-wide RNAi screens in bloodstream form trypanosomes identify drug transporters. Molecular & Biochemical Parasitology. 2011;175(1):91–4. doi:10.1016/j.molbiopara.2010.09.002

47. Malfara MF, Silverberg LJ, DiMaio J, Lagalante AF, Olsen MA, Madison E, et al. 2,3-Diphenyl-2,3-dihydro-4*H*-1,3-thiaza-4-one heterocycles inhibit growth and block completion of cytokinesis in kinetoplastid parasites. Molecular and Biochemical Parasitology. 2021 Sep 1;245:111396. doi:10.1016/j.molbiopara.2021.111396

48. Szöőr B, Simon DV, Rojas F, Young J, Robinson DR, Krüger T, et al. Positional Dynamics and Glycosomal Recruitment of Developmental Regulators during Trypanosome Differentiation. mBio. 2019 Jul 9;10(4):e00875–19. doi:10.1128/mBio.00875-19 PubMed PMID: 31289175; PubMed Central PMCID: PMC6747725.

49. Fleming J, Magana P, Nair S, Tsenkov M, Bertoni D, Pidruchna I, et al. AlphaFold Protein Structure Database and 3D-Beacons: New Data and Capabilities. Journal of Molecular Biology. 2025 Aug 1;Computation Resources for Molecular Biology437(15):168967. doi:10.1016/j.jmb.2025.168967

50. Mirdita M, Schütze K, Moriwaki Y, Heo L, Ovchinnikov S, Steinegger M. ColabFold: making protein folding accessible to all. Nat Methods. 2022 Jun;19(6):679–82. doi:10.1038/s41592-022-01488-1

51. Wheeler RJ. A resource for improved predictions of Trypanosoma and Leishmania protein three-dimensional structure. PLOS ONE. 2021 Nov 11;16(11):e0259871. doi:10.1371/journal.pone.0259871

52. Alvarez-Jarreta J, Amos B, Aurrecoechea C, Bah S, Barba M, Barreto A, et al. VEuPathDB: the eukaryotic pathogen, vector and host bioinformatics resource center in 2023. Nucleic Acids Res. 2024 Jan 5;52(D1):D808–16. doi:10.1093/nar/gkad1003

53. Pettersen EF, Goddard TD, Huang CC, Meng EC, Couch GS, Croll TI, et al. UCSF ChimeraX: Structure visualization for researchers, educators, and developers. Protein Science. 2021;30(1):70–82. doi:10.1002/pro.3943

54. Goddard TD, Huang CC, Meng EC, Pettersen EF, Couch GS, Morris JH, et al. UCSF ChimeraX: Meeting modern challenges in visualization and analysis. Protein Science. 2018;27(1):14–25. doi:10.1002/pro.3235

55. Meng EC, Goddard TD, Pettersen EF, Couch GS, Pearson ZJ, Morris JH, et al. UCSF ChimeraX: Tools for structure building and analysis. Protein Science. 2023;32(11):e4792. doi:10.1002/pro.4792

56. Urbaniak MD, Guther MLS, Ferguson MAJ. Comparative SILAC Proteomic Analysis of *Trypanosoma brucei* Bloodstream and Procyclic Lifecycle Stages. Li Z, editor. PLoS ONE. 2012;7(5):e36619–8. doi:10.1371/journal.pone.0036619

57. Nett IRE, Martin DMA, Miranda-Saavedra D, Lamont D, Barber JD, Mehlert A, et al. The Phosphoproteome of Bloodstream Form *Trypanosoma brucei*, Causative Agent of African Sleeping Sickness. Mol Cell Proteomics. 2009 Jul;8(7):1527–38. doi:10.1074/mcp.M800556-MCP200 PubMed PMID: 19346560; PubMed Central PMCID: PMC2716717.

58. Schimanski B, Nguyen TN, Günzl A. Highly efficient tandem affinity purification of trypanosome protein complexes based on a novel epitope combination. Eukaryotic Cell. 2005;4(11):1942–50. doi:10.1128/ec.4.11.1942-1950.2005

59. Vásquez-Trincado C, Patel M, Sivaramakrishnan A, Bekeová C, Anderson-Pullinger L, Wang N, et al. Adaptation of the heart to frataxin depletion: evidence that integrated stress response can predominate over mTORC1 activation. Hum Mol Genet. 2021 Sep 22;33(8):637–54. doi:10.1093/hmg/ddab216 PubMed PMID: 34550363; PubMed Central PMCID: PMC11000666.

60. Cox J, Mann M. MaxQuant enables high peptide identification rates, individualized p.p.b.-range mass accuracies and proteome-wide protein quantification. Nat Biotechnol. 2008 Dec;26(12):1367–72. doi:10.1038/nbt.1511 PubMed PMID: 19029910.

61. Liffner B, Absalon S. Expansion Microscopy Reveals Plasmodium falciparum Blood-Stage Parasites Undergo Anaphase with A Chromatin Bridge in the Absence of Mini-Chromosome Maintenance Complex Binding Protein. Microorganisms. 2021 Nov 6;9(11):2306. doi:10.3390/microorganisms9112306 PubMed PMID: 34835432; PubMed Central PMCID: PMC8620465.

62. Oliveira Souza RO, Jacobs KN, Back PS, Bradley PJ, Arrizabalaga G. IMC10 and LMF1 mediate mitochondrial morphology through mitochondrion-pellicle contact sites in Toxoplasma gondii. J Cell Sci. 2022 Nov 15;135(22):jcs260083. doi:10.1242/jcs.260083 PubMed PMID: 36314270; PubMed Central PMCID: PMC9845740.

63. Schindelin J, Arganda-Carreras I, Frise E, Kaynig V, Longair M, Pietzsch T, et al. Fiji: an open-source platform for biological-image analysis. Nat Methods. 2012 Jul;9(7):676–82. doi:10.1038/nmeth.2019

64. EPFL | Biomedical Imaging Group | TurboReg History [Internet]. [cited 2026 Mar 6]. Available from: https://bigwww.epfl.ch/publications/thevenaz1201.html

65. Thévenaz P, Ruttimann UE, Unser M. A pyramid approach to subpixel registration based on intensity. IEEE Trans Image Process. 1998;7(1):27–41. doi:10.1109/83.650848 PubMed PMID: 18267377.

66. Colasante C, Voncken F, Manful T, Ruppert T, Tielens AGM, van Hellemond JJ, et al. Proteins and lipids of glycosomal membranes from Leishmania tarentolae and Trypanosoma brucei. F1000Res. 2013 Jan 29;2:27. doi:10.12688/f1000research.2-27.v1 PubMed PMID: 24358884; PubMed Central PMCID: PMC3814921.

67. Hoepfner D, van den Berg M, Philippsen P, Tabak HF, Hettema EH. A role for Vps1p, actin, and the Myo2p motor in peroxisome abundance and inheritance in Saccharomyces cerevisiae. J Cell Biol. 2001 Dec 10;155(6):979–90. doi:10.1083/jcb.200107028 PubMed PMID: 11733545; PubMed Central PMCID: PMC2150915.

68. de Rooij IIS, Allwood EG, Aghamohammadzadeh S, Hettema EH, Goldberg MW, Ayscough KR. A role for the dynamin-like protein Vps1 during endocytosis in yeast. J Cell Sci. 2010 Oct 15;123(20):3496–506. doi:10.1242/jcs.070508

69. Koirala S, Guo Q, Kalia R, Bui HT, Eckert DM, Frost A, et al. Interchangeable adaptors regulate mitochondrial dynamin assembly for membrane scission. Proceedings of the National Academy of Sciences. 2013 Apr 9;110(15):E1342–51. doi:10.1073/pnas.1300855110

70. Lackner LL, Horner JS, Nunnari J. Mechanistic analysis of a dynamin effector. Science. 2009 Aug 14;325(5942):874–7. doi:10.1126/science.1176921 PubMed PMID: 19679814; PubMed Central PMCID: PMC6546417.

71. Williams C, Opalinski L, Landgraf C, Costello J, Schrader M, Krikken AM, et al. The membrane remodeling protein Pex11p activates the GTPase Dnm1p during peroxisomal fission. Proceedings of the National Academy of Sciences. 2015 May 19;112(20):6377–82. doi:10.1073/pnas.1418736112

72. Koch J, Pranjic K, Huber A, Ellinger A, Hartig A, Kragler F, et al. PEX11 family members are membrane elongation factors that coordinate peroxisome proliferation and maintenance. J Cell Sci. 2010 Oct 1;123(19):3389–400. doi:10.1242/jcs.064907

73. Schrader M, Reuber BE, Morrell JC, Jimenez-Sanchez G, Obie C, Stroh TA, et al. Expression of *PEX11*β Mediates Peroxisome Proliferation in the Absence of Extracellular Stimuli*. Journal of Biological Chemistry. 1998 Nov 6;273(45):29607–14. doi:10.1074/jbc.273.45.29607

74. Cayla M, McDonald L, MacGregor P, Matthews K. An atypical DYRK kinase connects quorum-sensing with posttranscriptional gene regulation in Trypanosoma brucei. Engstler M, Soldati-Favre D, Acosta Serrano Á, editors. eLife. 2020 Mar 26;9:e51620. doi:10.7554/eLife.51620

75. Benz C, Urbaniak MD. Organising the cell cycle in the absence of transcriptional control: Dynamic phosphorylation co-ordinates the Trypanosoma brucei cell cycle post-transcriptionally. PLOS Pathogens. 2019 Dec 12;15(12):e1008129. doi:10.1371/journal.ppat.1008129

76. Ford MGJ, Chappie JS. The Structural Biology of the Dynamin-Related Proteins: New Insights into a Diverse, Multi-Talented Family. Traffic. 2019 Oct;20(10):717–40. doi:10.1111/tra.12676 PubMed PMID: 31298797; PubMed Central PMCID: PMC6876869.

77. Voncken F, van Hellemond JJ, Pfisterer I, Maier A, Hillmer S, Clayton C. Depletion of GIM5 Causes Cellular Fragility, a Decreased Glycosome Number, and Reduced Levels of Ether-linked Phospholipids in Trypanosomes *. Journal of Biological Chemistry. 2003 Sep 12;278(37):35299–310. doi:10.1074/jbc.M301811200 PubMed PMID: 12829709.

78. Ekal L, Alqahtani AMS, Hettema EH. The dynamin-related protein Vps1 and the peroxisomal membrane protein Pex27 function together during peroxisome fission. J Cell Sci. 2023 Mar 24;136(6):jcs246348. doi:10.1242/jcs.246348

79. Schrader TA, Carmichael RE, Islinger M, Costello JL, Hacker C, Bonekamp NA, et al. PEX11β and FIS1 cooperate in peroxisome division independently of mitochondrial fission factor. J Cell Sci. 2022 Jul 8;135(13):jcs259924. doi:10.1242/jcs.259924

80. Imoto Y, Itoh K, Fujiki Y. Molecular Basis of Mitochondrial and Peroxisomal Division Machineries. International Journal of Molecular Sciences. 2020 Jul 29;21(15). doi:10.3390/ijms21155452

81. Schrader M. Shared components of mitochondrial and peroxisomal division. Biochimica et Biophysica Acta (BBA) - Molecular Cell Research. 2006 May 1;Mitochondrial Dynamics in Cell Life and Death1763(5):531–41. doi:10.1016/j.bbamcr.2006.01.004

82. Vertommen D, Van Roy J, Szikora JP, Rider MH, Michels PAM, Opperdoes FR. Differential expression of glycosomal and mitochondrial proteins in the two major life-cycle stages of *Trypanosoma brucei*. Molecular and Biochemical Parasitology. 2008 Apr 1;158(2):189–201. doi:10.1016/j.molbiopara.2007.12.008

83. Iyer A, Hemphill A, Schneider A. TbDLP and TbFis1 mediate glycosomal morphology and biogenesis in Trypanosoma brucei [SSRN Scholarly Paper] [Internet]. Rochester, NY: Social Science Research Network; 2025 [cited 2026 Mar 8]. Available from: https://papers.ssrn.com/abstract=5744657 doi:10.2139/ssrn.5744657

84. Bauer S, Morris MT. Glycosome biogenesis in trypanosomes and the de novo dilemma. PLOS Neglected Tropical Diseases. 2017 Apr 20;11(4):e0005333. doi:10.1371/journal.pntd.0005333

85. Sugiura A, Mattie S, Prudent J, McBride HM. Newly born peroxisomes are a hybrid of mitochondrial and ER-derived pre-peroxisomes. Nature. 2017 Feb;542(7640):251–4. doi:10.1038/nature21375

86. Benz C, Müller N, Kaltenbrunner S, Váchová H, Vancová M, Lukeš J, et al. Kinetoplastid-specific X2-family kinesins interact with a kinesin-like pleckstrin homology domain protein that localizes to the trypanosomal microtubule quartet. Molecular Microbiology. 2022;118(3):155–74. doi:10.1111/mmi.14958

87. Hilton NA, Sladewski TE, Perry JA, Pataki Z, Sinclair-Davis AN, Muniz RS, et al. Identification of TOEFAZ1-interacting proteins reveals key regulators of *Trypanosoma brucei* cytokinesis: *TOEFAZ1 interactors in* T. brucei. Molecular Microbiology. 2018 Aug;109(3):306–26. doi:10.1111/mmi.13986

